# An optogenetic method for interrogating YAP1 and TAZ nuclear-cytoplasmic shuttling

**DOI:** 10.1101/2020.06.08.140228

**Authors:** Anna M. Dowbaj, Robert P. Jenkins, John M. Heddleston, Alessandro Ciccarelli, Todd Fallesen, Klaus Hahn, Marco Montagner, Erik Sahai

**Author notes:** these authors contributed equally (Anna M. Dowbaj, Robert P. Jenkins).

## Abstract

The shuttling of transcription factors and transcriptional regulators in and out of the nucleus is central to the regulation of many biological processes. Here we describe a new method for studying the rates of nuclear entry and exit of transcriptional regulators. A photo-responsive AsLOV (*Avena sativa* Light Oxygen Voltage) domain is used to sequester fluorescently-labelled transcriptional regulators YAP1 and TAZ/WWTR1 on the surface of mitochondria and reversibly release them upon blue light illumination. After dissociation, fluorescent signals from mitochondria, cytoplasm and nucleus are extracted with a bespoke app and used to generate rates of nuclear entry and exit. Using this method, we demonstrate that phosphorylation of YAP1 on canonical sites enhances its rate of nuclear export. Moreover, we provide evidences that, despite high intercellular variability, YAP1 import and export rates correlated within the same cell. By simultaneously releasing YAP1 and TAZ from sequestration, we show that their rates of entry and exit are correlated. Furthermore, tracking of light-sensitive YAP1 with lattice lightsheet microscopy revealed high heterogeneity of YAP1 dynamics within different subcellular regions, suggesting that implementing high resolution volumetric 3D data could shed light on new mechanisms of nuclear-cytoplasmic shuttling of proteins.

## Introduction

Rapid regulation of cellular processes in space and time is mostly achieved either by switching on inactive proteins in the correct location, or by recruiting proteins at the appropriate subcellular compartment at the right time. Compartmentalisation is a fundamental feature of eukaryotic cells and numerous membrane bound organelles contribute to sub-cellular organisation. Shuttling of proteins between compartments is a key mechanism of regulating many processes, including the transcription of DNA in the nucleus (Fu et al., 2018, Xu and Massague, 2004, Di Ventura and Kuhlman, 2016). Transcription factors and transcriptional regulators are examples of proteins regulated by compartmentalisation and their nuclear accumulation is often restricted in absence of activating signals, either by cytoplasmic tethering or constitutive nuclear export (Xu and Massague, 2004). Mechanistically, members of the karyopherin family recognize specific signals on cargo proteins (larger than 40KDa), namely nuclear localization signals (NLS) and nuclear export signals (NES), and mediate their transition across the nuclear pore complex (NPC), while loading and unloading of cargoes are controlled by the GTPase Ran (Fu et al., 2018, Di Ventura and Kuhlman, 2016), thus representing an energetic barrier to NC shuttling. Importantly, the function of many transcription factors and regulators is the integration of several signaling inputs. The specificity of the response is dictated by the modulation of frequency, intensity and duration of nuclear-cytoplasmic (NC) shuttling rather than linear nuclear accumulation (Hao and O’Shea, 2011, Purvis and Lahav, 2013, Gao et al., 2018, Yosef and Regev, 2011, Behar and Hoffmann, 2010). This change in our understanding highlights the importance of live, non-destructive techniques.

YAP1 and TAZ transcriptional regulators are a perfect example of such complexity. They serve as a hub for a wide range of stimuli, including biochemical (Hippo, (Meng et al., 2016)) and metabolic pathways (Sorrentino et al., 2014, Bertolio et al., 2019), mechanical inputs (rigidity, shape, stiffness, cell density and shear stress) (Wada et al., 2011, Calvo et al., 2013, Benham-Pyle et al., 2015, Aragona et al., 2013, Dupont et al., 2011, Chaudhuri et al., 2016, Nakajima et al., 2017, Elosegui-Artola et al., 2017), polarity and other signalling cascades (GPCRs, Akt, JNK) (Calvo et al., 2013, Codelia et al., 2014, Basu et al., 2003) (Fig. S1A). YAP1 and TAZ are master regulators of cell proliferation, apoptosis, phenotypic plasticity (Zanconato et al., 2019, Shreberk-Shaked and Oren, 2019) and unsurprisingly they are involved in all the steps of tumorigenesis (Zanconato et al., 2016) (Fig. S1A). In the canonical view of the pathway, YAP1 and TAZ activity is inhibited by nuclear exclusion upon LATS1/2 (the main kinases of Hippo pathway downstream of MST1/2) phosphorylation on key serine residues, and subsequent binding to 14-3-3 proteins or proteasomal degradation. When the phosphorylation is low, YAP1 and TAZ can accumulate into the nucleus and bind to their DNA-binding partners TEAD1-4, BRD4 and MED1, which also contribute to their nuclear accumulation. However, this over simplistic model has been challenged by recent findings that LATS-phosphorylated-YAP1 can be found in the nucleus and that nuclear accumulation is not sufficient for YAP1-driven transcriptional activity (Ege et al., 2018, Wada et al., 2011). Moreover, several LATS-independent mechanisms have been shown to affect YAP1 and TAZ NC shuttling and activity, such as nuclear deformation (Elosegui-Artola et al., 2017), cell cycle (Manning et al., 2018), Src family kinases (Calvo et al., 2013, Ege et al., 2018, Elbediwy et al., 2016, Rosenbluh et al., 2012, Tamm et al., 2011) and, more recently, phase separation (Lu et al., 2020, Cai et al., 2019). Despite the intense studies on these transcription regulators during the years, there is no consensus on how their NC shuttling is regulated.

Historically NC shuttling has been investigated with static methods, on fixed cells or subcellular fractions, with low temporal and spatial resolution, and using chemical such as the export inhibitor Leptomycin B. More recently, measurement of protein shuttling has been revolutionised by the use of fluorescent dyes and proteins (Molenaar and Weeks, 2018, Di Ventura and Kuhlman, 2016). Without perturbation, the partitioning of fluorescent signal between compartment reflects the equilibrium position of the various shuttling process influencing the fluorescently tagged protein. The application of light-driven perturbations, such as photobleaching, provides more information about rates of transit between compartments. However, one downside of photobleaching is that it is inherently destructive by irreversibly damaging the fluorophore, which limits the researcher’s ability to make repeated measurements and typically generates only a single intensity decay curve, which limits the mathematical fitting to derive shuttling rates.

During the last decade, several optogenetic systems have been developed to control NC localisation of proteins within a few seconds upon light illumination. Early examples of light-mediated NC shuttling (Engelke et al., 2014, Crefcoeur et al., 2013) led to irreversible nuclear accumulation and they have been surpassed by reversible methods based on light-sensitive proteins. The common idea underlying optogenetic systems is the light-dependent conformational change of photoreceptors fused with the proteins of interest, that can be exploited to manipulate a wide range of cellular processes (de Mena et al., 2018). Desired protein localization upon light illumination has been achieved with two strategies so far: two-components systems, in which a bait protein is anchored to one compartment and the interaction with the prey protein is light-dependent (Guntas et al., 2015, Kennedy et al., 2010, Beyer et al., 2015, Strickland et al., 2012), and one-component systems, in which NES/NLS peptides are caged/exposed upon light illumination (Niopek et al., 2014, Niopek et al., 2016, Yumerefendi et al., 2015, Yumerefendi et al., 2016). Interestingly, the former strategy has been recently used to develop light-induced control of cellular forces and, as a consequence, of YAP1 localisation (Valon et al., 2017). In this work, we exploited an optogenetic tool, namely LOV-TRAP (Wang and Hahn, 2016, Wang et al., 2016a), for studying NC shuttling based upon light-induced release and sequestration of the protein of interest (POI). We present detailed methods for the molecular biology, imaging, and analysis of this tool. Furthermore, the utility of the method is demonstrated by measuring the nuclear import and export rates of mCherry, YAP1, YAP1 mutants, and TAZ. We show that import and export rates, despite the wide intercellular variability, are correlated within the same cell, and that the export rate is the main determinant of wild-type YAP1 distribution. Finally, we provide the proof of principle that multiple proteins can be recorded simultaneously within the same cell by analysing YAP1 and TAZ nuclear transit rates.

## Results

To establish an optogenetic system for protein shuttling studies, we adapted the LOV-TRAP system (Wang and Hahn, 2016, Wang et al., 2016a). With this tool, a mitochondria-tethered LOV domain interacts with its binding partner Zdk98 (Zdk hereafter), fused to the POI labelled with a fluorescent marker. Upon blue-light illumination, LOV and Zdk dissociate, releasing the POI into the cytoplasm within a fraction of a second and enabling subsequent monitoring. We selected the outer surface of mitochondria for sequestration because their distinctive and discrete morphology is well suited to image segmentation. Furthermore, the outer membrane is directly in contact with a large area of the cytoplasm. Therefore, we fused the LOV domain to the mitochondrial anchor sequence TOM20, while the POI was fused to a Zdk peptide sequence. The fluorophore mCherry was selected to visualise the target protein because it is excited with sufficiently long wavelength light to exclude the possibility of triggering a conformational change in the LOV domain. Fig. 1A and Fig. S1B shows a schematic diagram of the constructs and their expected localisation, and expected response to blue light inducing a conformational change in the LOV domain.

**Figure 1:**
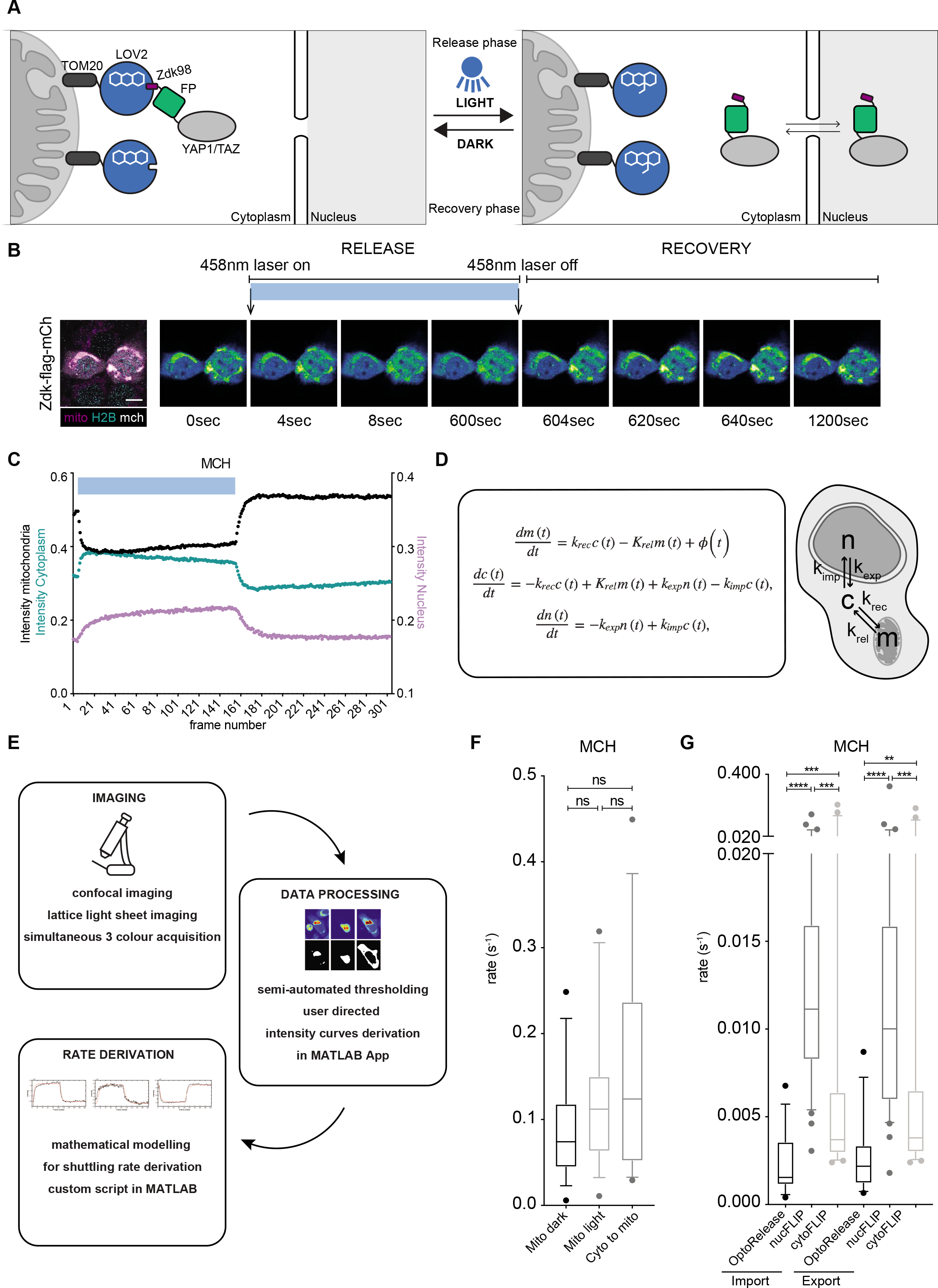
Opto-release system. **(A)** Opto-release system specifications; In the dark, Zdk-tagged protein is tightly sequestered on the mitochondria through the interaction with light sensitive LOV domain. The LOV domain is tethered to the mitochondria through TOM20 protein anchor. Upon blue light illumination, the LOV domain changes conformation and releases Zdk-tagged protein. **(B)** Representative image of Zdk-flag-mCh constructs release and recovery from mitochondrial sequestration by TOM20-flag-LOVwt. Experiment with 10 min of blue light illumination (the release phase), followed by 10 min of dark recovery without blue light (the recovery phase). Scale bar is 10μm. **(C)** Quantification of intensities corresponding to optogenetic release and recovery experiment performed in HaCaT cells transiently transfected with Zdk-flag-mCherry (n=15 cells). Only the mean of all cells from at least 3 biological replicates is shown for clarity. Intensity has been corrected for bleaching and background signal. **(D)** System of ordinary differential equations used to derive the shuttling rates of import, *k_imp_*, export, *k_exp_*, rate of mitochondrial release, *K_rel_*, and rate of mitochondrial recovery, *k_rec_* for the nucleus, *n*(*t*), cytoplasm, *c*(*t*) and mitochondria, *m*(*t*) at time *t*. The function *ϕ*(*t*) allows the fitting of the escape of intensity to elsewhere in the cell. **(E)** Experimental pipeline. Confocal imaging is followed by semi-automated intensity thresholding and shuttling rate analysis with mathematical modelling. **(F)** The numerical values of mitochondrial sequestration rates at steady state (darkness), during release and during recovery phases, compared to nuclear import/export rates for Zdk-flag-mCherry (n=16), Number of cells is given in brackets. Boxplot (10&90 percentile, median) represents at least three independent biological replicates. Mann-Whitney U test. **(G)** Zdk-flag-mCherry nuclear import and export rates acquired using nuc/cyto FLIP method compared to the optogenetics method. Boxplot (10&90 percentile, median) represents at least three independent biological replicates. The values of optogenetic-derived import/export rates from G are replotted here for clarity and comparison to FLIP. Mann-Whitney U test.

### An opto-release system for studying nuclear cytoplasmic shuttling

We began by testing if a fusion of the Zdk peptide to mCherry was capable of localising the fluorescent protein to mitochondria in the presence of the TOM20-LOV protein. Cells expressing histone H2B-mTurquoise fusion were co-transfected with these constructs and additionally labelled with a MitoTracker™ dye to allow visualisation of the different compartments. Fig. 1B shows that this approach labels the nucleus and mitochondria in the expected way. Importantly, Zdk-mCherry exhibited expected enrichment to mitochondria in the presence of the TOM20-LOV fusion (Fig. 1B and Fig. S1C, where we show localisation without sequestration). All the constructs used in this study have expected localisation, size and transcriptional activity without sequestration, confirming that these parameters were not affected by fusion with Zdk and mCherry (Fig. S1D-F). We next investigated the effect of 458 nm light illumination. Fig. 1B shows that blue light illumination led to reduced mitochondrial localisation and increased Zdk-mCherry in both the cytoplasm and nucleus. Upon cessation of illumination, the Zdk-mCherry fusion exhibited a progressive transition back to the mitochondria. Quantification of the fluorescent signal in mitochondrial, cytoplasmic, and nuclear compartments confirmed the release and re-binding of Zdk-mCherry to the mitochondria, illustrated in Fig. 1C. Of note, the rate of fluorescence increase and decrease in the nucleus was much slower than that in the cytoplasm. This is most likely related to the rate of nuclear entry and exit of Zdk-mCherry being slower that the kinetics of release and re-binding of the Zdk peptide to the LOV domain. We reasoned that this difference could be exploited to gain quantitative information about the rates of nuclear import and export of proteins. To this end, we developed a computational tool for automatically segmenting the mitochondria, nuclear, and peri-nuclear cytoplasm excluding mitochondria. The time series data generated for each compartment (Fig. 1C) was then fitted to a series of ordinary differential equations (ODEs) (Fig. 1D), with a workflow demonstrated in Fig. 1E.

### Supervised semi-automated segmentation of cellular compartments using a MATLAB app

To accurately extract quantitative information about the localisation of the fluorescent Zdk-fusion, we developed a supervised MATLAB app for thresholding the different cellular compartments. Upon loading the imaging data and defining the appropriate channels (Fig. S2A), the region containing the cell of interest can be selected. Percentile intensity projections of all frames in each relevant channel are made at levels appropriate to bring out the features of each compartment whilst minimising noise and interference from neighbouring cells (Fig S2A). The percentile projections are then thresholded to define appropriate compartment boundaries. The appropriate threshold for identifying the whole cell is then set using a sliding tool that provides a real-time image of the result of the thresholding (Fig. S2C). Once an appropriate value has been selected, the process is repeated for the thresholding of the mitochondria, and then the nucleus (Fig. S2D). In the analysis presented here, nuclear segmentation was based on the fluorescent histone image and mitochondrial segmentation was based on MitoTracker™. Thresholding of the cytoplasmic compartment was used to define a region around the nucleus, with mitochondrial regions automatically excluded. We typically chose a threshold that excluded thinner parts of the cell periphery that did not fill the whole z-plane being acquired. Once thresholds are set, the app generates both visual plots and a numerical data file to allow model fitting and the estimation of rates of binding and unbinding of the fluorescent Zdk fusion protein from mitochondria and its rates of nuclear entry and exit (Fig. S2D). To account for possible changes in the intensity of fluorescent cellular, nuclear, and mitochondrial labels during the imaging, the thresholding parameters could be re-defined at varying points during the time series. The result of this dynamic thresholding could then be compared with constant threshold values, with the method giving smaller errors in fitting selected for subsequent quantitative analysis.

### Differential equations to model nuclear import and export

Having generated numerical data of the localisation of Zdk-fusion protein of interest over time, we then sought to determine the rates of nuclear entry and exit. We used ODEs to model this as a system with three key variables: the amount of Zdk-fusion bound to mitochondria, the amount in the cytoplasm, and the amount in the nucleus (Fig. 1D). Transitions were permitted from the mitochondria to the cytoplasm, and vice-versa, and from the cytoplasm to the nucleus, and vice-versa. The rate of transition on and off mitochondria and rates from cytoplasm to nucleus and nucleus to cytoplasm transfer were fitted simultaneously from the system of ODEs. To account for any photobleaching, we applied a normalisation method to ensure that the total level of fluorescence was constant over time. This improved the data fitting.

Fig. 1F,G show the directly measured mitochondrial release and re-binding rates and the inferred rates of Zdk-mCherry nuclear entry and exit. As we expected, the rates of transit on and off the mitochondria were considerably faster than the nuclear transit rates (compare Fig. 1F with “Optorelease” samples in Fig. 1G). Our methodology uses information from both the release and re-binding phases of the experiment to determine the nuclear entry and exit rates. Fig. S3A,B shows the advantage that this confers over using only the release or only the re-binding phase, with two-fold and eight-fold reductions in the intervals of plausible fitted parameter values when both release and re-binding phases are used compared to release and re-binding phases, respectively. We sought to corroborate the rates of nuclear entry and exit using a conventional fluorescence loss in photobleaching method (FLIP). In this method, we continuously deplete the Zdk-fluorescence at a location either in the cytoplasm (cytoFLIP) or the nucleus (nucFLIP) and record the diffusion of bleached fluorophore in the other compartment. Fig. 1G shows good concordance between the rates measured using our opto-release method and cytoFLIP. Of note, if we performed nucFLIP with continuous bleaching at a point in the nucleus, then the rates of nuclear entry and exit diverged from those measured using both cytoFLIP and opto-release. More specifically, the inferred rates were faster. This is most probably due to direct photobleaching of the fluorophore in the cytoplasm above and below the nucleus. This highlights an important caveat of FLIP experiments – namely the choice of photobleaching location.

### Application of opto-release methodology to YAP1 and TAZ

Having established that our method involving light-dependent release of a molecule from sequestration on the outer surface of mitochondria was capable of measuring nuclear entry and exit rates, we applied our method to measure the nuclear import and export rates of the transcriptional regulators YAP1 and TAZ. YAP1 and TAZ were fused to the Zdk peptide and mCherry. Fig. S1E confirms that these chimeric proteins retained their expected ability to drive transcription from a TEAD-dependent promoter, indicating that fusion with fluorophore and Zdk peptide didn’t affect binding to other proteins. As before, blue light was used to release YAP1 and TAZ from sequestration and their redistribution into the cytoplasm and nucleus was tracked. After 600 seconds, the blue light illumination was stopped and the return of signal to the mitochondria was monitored (Fig. 2A). The app interface described above was then used to extract quantitation (Fig. 2B,C) that was then fitted to the model (Fig. 2D-H). These analyses reveal that both YAP1 and TAZ have slightly lower mitochondria release/recovery rates (Fig. 2D-F), but elevated nuclear import and export rates compared to mCherry, with TAZ rates being slightly faster than the rates of transit across the nuclear envelope for YAP1 (Fig. 2G,H). Once again, we confirmed that measurements obtained by optogenetic release and re-sequestration of YAP1 agreed with data obtained using more conventional FLIP methodology (Fig. S4A-I).

**Figure 2:**
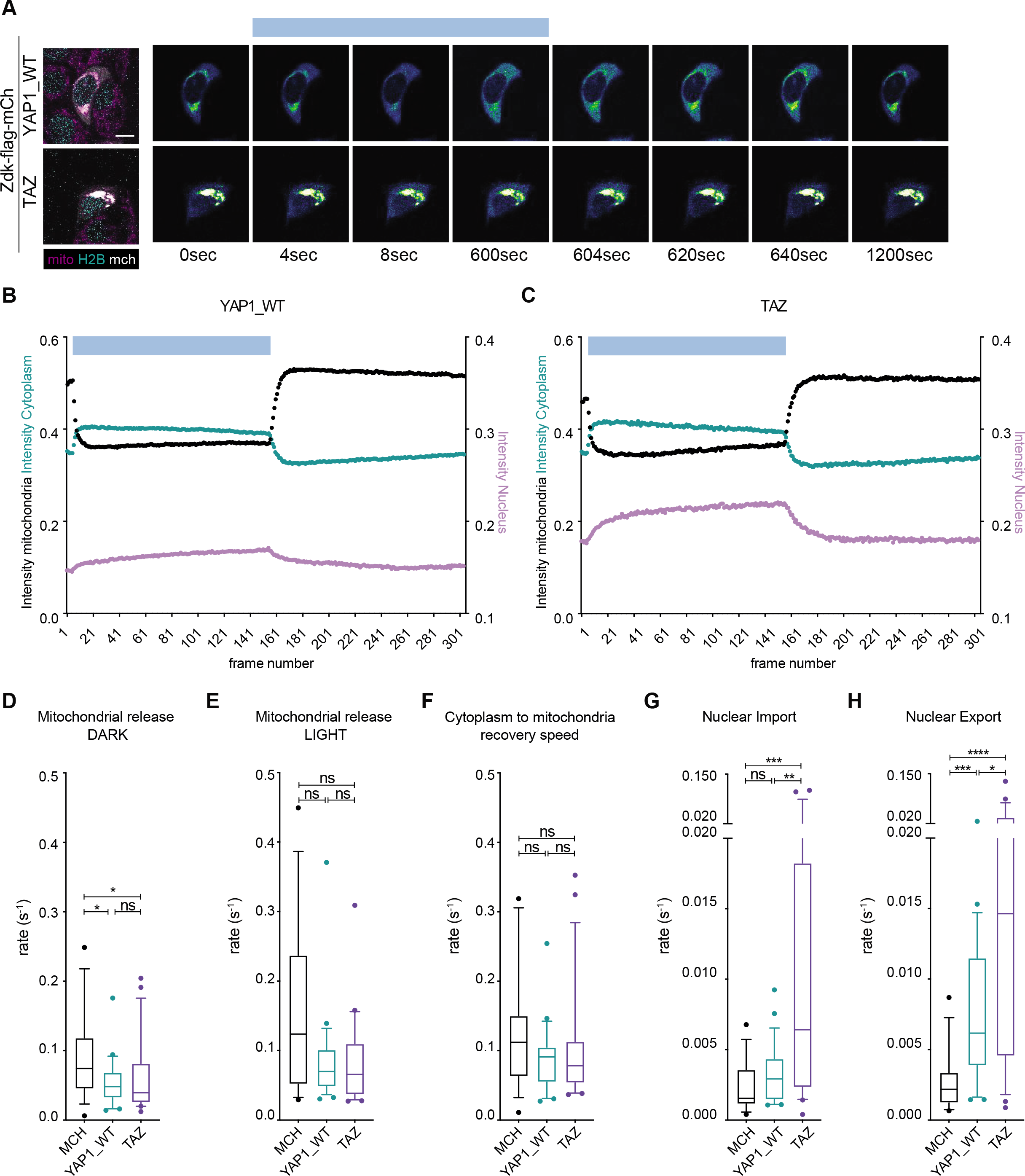
Rates of YAP1 and TAZ nucleocytoplasmic shuttling in HaCaT epithelial cell line. **(A)** Representative images of Zdk-flag-mCh-YAP1_WT/TAZ constructs release and recovery from mitochondrial sequestration by TOM20-flag-LOVwt. Experiment with 10 min of blue light illumination (the release phase), followed by 10 min of dark recovery without blue light (the recovery phase). Scale bar is 10μm. **(B) (C)** Quantification of absolute intensities corresponding to optogenetic release and recovery experiment performed in HaCaT cells transiently transfected with (B) Zdk-flag-mCh-YAP1_WT (n=24) or (C) Zdk-flag-mCh-TAZ (n=22). Only the mean of all cells from at least 3 biological replicates is shown for clarity. Intensity has been corrected for bleaching and background signal. **(D)(E)(F)** The numerical values of mitochondrial sequestration rates for Zdk-flag-mCherry-YAP1_WT (n=28) and Zdk-flag-mCh-TAZ (n=22) in steady state (darkness), during release and during recovery phases. Number of cells per each condition is given in brackets. Boxplot (10&90 percentile, median) represents at least three independent biological replicates. Mann-Whitney U test. **(G)(H)** YAP1 and TAZ nuclear import and export rates. Boxplot (10&90 percentile, median) represents at least three independent biological replicates. Mann-Whitney U test.

### Active YAP1 is subject to reduced rates of nuclear export

Compartmentalization of YAP1 and TAZ is regulated by different post-translational modifications and, despite recent evidences on the importance of methylation and acetylation, the most characterised is phosphorylation. YAP1 and TAZ are negatively regulated by LATS1/2-dependent phosphorylations, downstream of the Hippo pathway, and conversion of these residues to alanine generates a LATS1/2-insensitive YAP1_5SA mutant that shows nuclear localisation and higher transcriptional activity. On the contrary, the YAP1_S94A mutant is transcriptionally inactive and accumulates in the cytoplasm, due to its inability to bind the transcriptional cofactor and nuclear tether TEAD. We investigated if YAP1_5SA and YAP1_S94A mutants exhibited different nuclear import and export rates. Fig. 3A-C and Fig. S5A-E show that the most prominent effect of 5SA mutations is reduced nuclear export compared to the wild-type isoform. In contrast, this doesn’t hold true for the YAP1_S94A mutant, that shows rates comparable to the wild-type isoform. Rates of nuclear import and export are consistently correlated as shown for YAP1 and its mutants in Fig. 3C. Cells with high rates of YAP1 nuclear import also had high rates of nuclear export. These data suggest that the energetic cost of transit across the nuclear pore might be highly variable between cells, but that this variation in cost applies to both nuclear entry and exit. Furthermore, even though there is considerable inter-cellular variation in the rates, the process remains subject to regulation. These results support our previous findings that YAP1_5SA nuclear persistence can be explained by slower nuclear exit, while cytoplasmic enrichment of S94A at steady state cannot simply explained by regulation of NC shuttling and thus requires additional explanations, such as tethering by cytoplasmic proteins (Ege et al., 2018).

**Figure 3:**
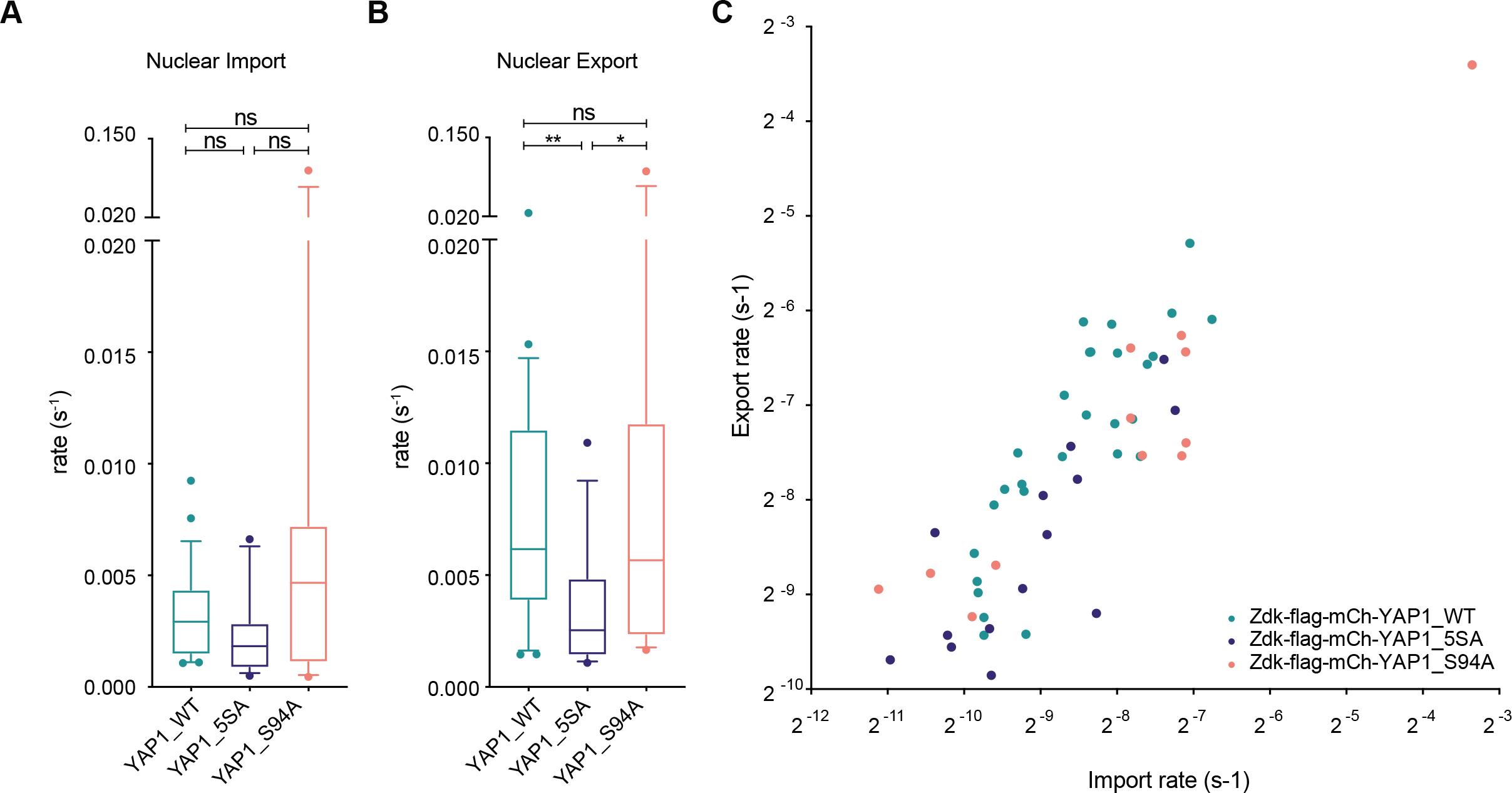
Rates of YAP1 mutants nucleocytoplasmic shuttling in epithelial cell line. **(A)(B)** YAP1 mutants nuclear import and export rates. Boxplot (10&90 percentile, median) represents at least three independent biological replicates. Mann-Whitney U test. **(C)** Scatterplot of import versus export and line of the best fit for Zdk-flag-mCherry-YAP1_WT (teal), Zdk-flag-mCherry-YAP1_5SA (dark blue), Zdk-flag-mCherry-YAP1_S94A (orange).

### Import and export rates are correlated

We were intrigued that the variance in import and export rates spanned an order of magnitude and investigated if this variation might correlate with differences in cell or nuclear morphology, as has been suggested for nuclear import previously (Elosegui-Artola et al., 2017). Fig. 4A-E shows correlation analysis of import and export rates with a range of nuclear morphology parameters. Some correlations are trivial, such as the positive association between area and perimeter, whereas as others are more intriguing, for example, the ratio of import and export rates was strongly correlated with the relative NC distribution. This latter metric reflects the relative segregation of any molecules that were not effectively sequestered by the LOV domain prior to the period of illumination. This is reassuring because in a simple model the ratio between relative rates of import and export should reflect the distribution of molecules between the two compartments at steady state. However, it is not possible to attach too much significance to this as the initial nuclear and cytoplasmic distributions of the fluorophore are assumed to be linearly proportional to the fitted compartment transfer rates and initial mitochondrial intensity in the model fitting. Thus, this observation reflects that our modelling approach is internally consistent. That said, import/export ratio positively correlates with NC distribution for all the proteins analysed, whereas single values of import and export did not, suggesting a biological information beyond the mere technical control. Similarly, import/export ratio positively correlates with nuclear area to cytoplasmic area ratio. Since fitting population intensities, this might reflect that larger nuclei will contain relatively greater levels of fluorescent molecules. Interestingly, the cytoplasmic shape parameters did not show any significant correlations with import and export rates (Fig. S6A-E). More intriguingly, we note that only in the case of YAP1 there was a negative correlation between the export rate and nuclear/cytoplasmic ratio (Fig. 4F). This suggests that tuning of the export rate is the main determinant of the overall nuclear/cytoplasmic distribution of YAP1 and that it becomes reduced as the nucleus takes up a larger area of the cell. Plots relating nuclear import to the same parameters did not show any correlation. The lack of correlation between export rate of mCherry, TAZ, or the YAP1 mutants, with nuclear/cytoplasm ratio or nuclear area/cytoplasmic area indicates that this regulation is specific to wild-type YAP1 and that some mechanisms don’t apply to YAP1 mutants (such as cytoskeletal regulation of YAP1_5SA mutant (Ege et al., 2018).

**Figure 4:**
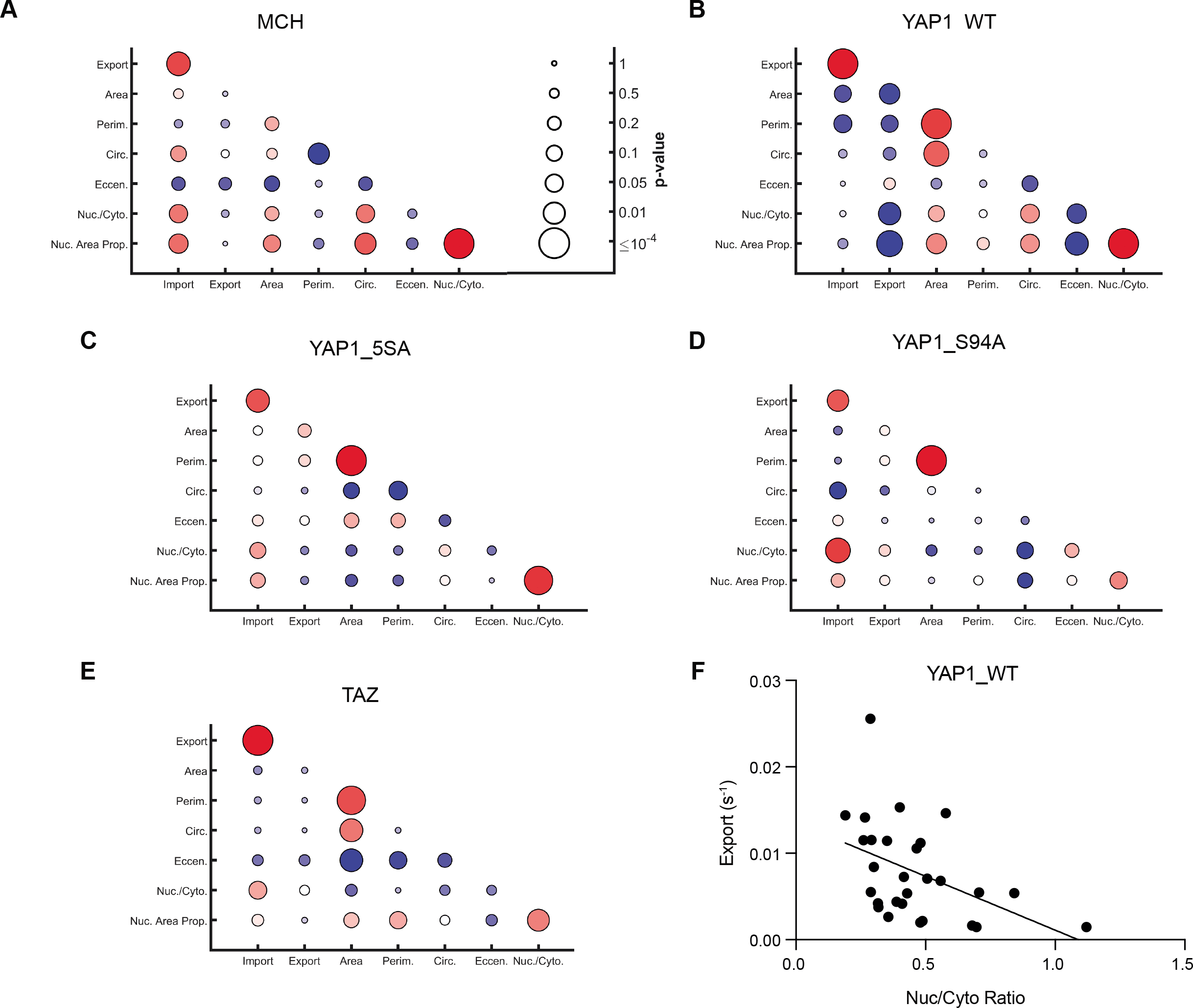
Nuclear morphology correlation with import and export. **(A)(B)(C)(D)(E)** Import and export correlations with nuclear morphology, specified by area, perimeter, circularity and eccentricity, as well as nuclear-to-cytoplasmic ratio and nuclear area as proportion of cell. Plots show Zdk-flag-mCherry (A), Zdk-flag-mCh-YAP1_WT (B), Zdk-flag-mCh-YAP1_5SA (C), Zdk-flag-mCh-YAP1_S94A (D) and Zdk-flag-mCh-TAZ (E). Circle colour reflects Pearson correlation (bright red +1, dark blue −1) and circle size the p-value of the correlation (large, significant; small, nonsignificant). **(F)** Negative correlation between export and NC ratio of YAP1_WT.

### Simultaneous measurement of YAP1 and TAZ nuclear transit rates reveals that they are correlated

The observation of variance in nuclear transit rates between cells prompted us to wonder if the transit rates of two different proteins in the same cell might be correlated. Testing this idea using conventional FLIP methodology would be highly challenging as it would require both bleaching and imaging two fluorophores simultaneously, with significant potential for unintended bleaching artefacts. However, using the opto-release system with two differently labelled POIs would have fewer problems. Therefore, we labelled YAP1 with Venus and TAZ with mCherry and confirmed that proteins could be tracked in parallel (Fig. 5A). Fig. 5B,C and Fig. S7A-C show that mitochondrial release and nuclear import are similar between YAP1 and TAZ, but the latter shows increased nuclear export (Fig. 5C). This suggests a lower tethering of TAZ in the nucleus compared to YAP1, maybe due to the relative contribution of different mechanisms, such as YAP1 regulation by the cytoskeleton (Dupont et al., 2011, Calvo et al., 2013) and the presence of different NLS and NES sequences in YAP1 and TAZ (Kofler et al., 2018, Fang et al., 2018). Importantly, both export and import rates of YAP1 significantly correlate with those of TAZ, indicating still a large overlap between the mechanisms regulating NC shuttling of the two proteins (Fig. 5D,E).

**Figure 5:**
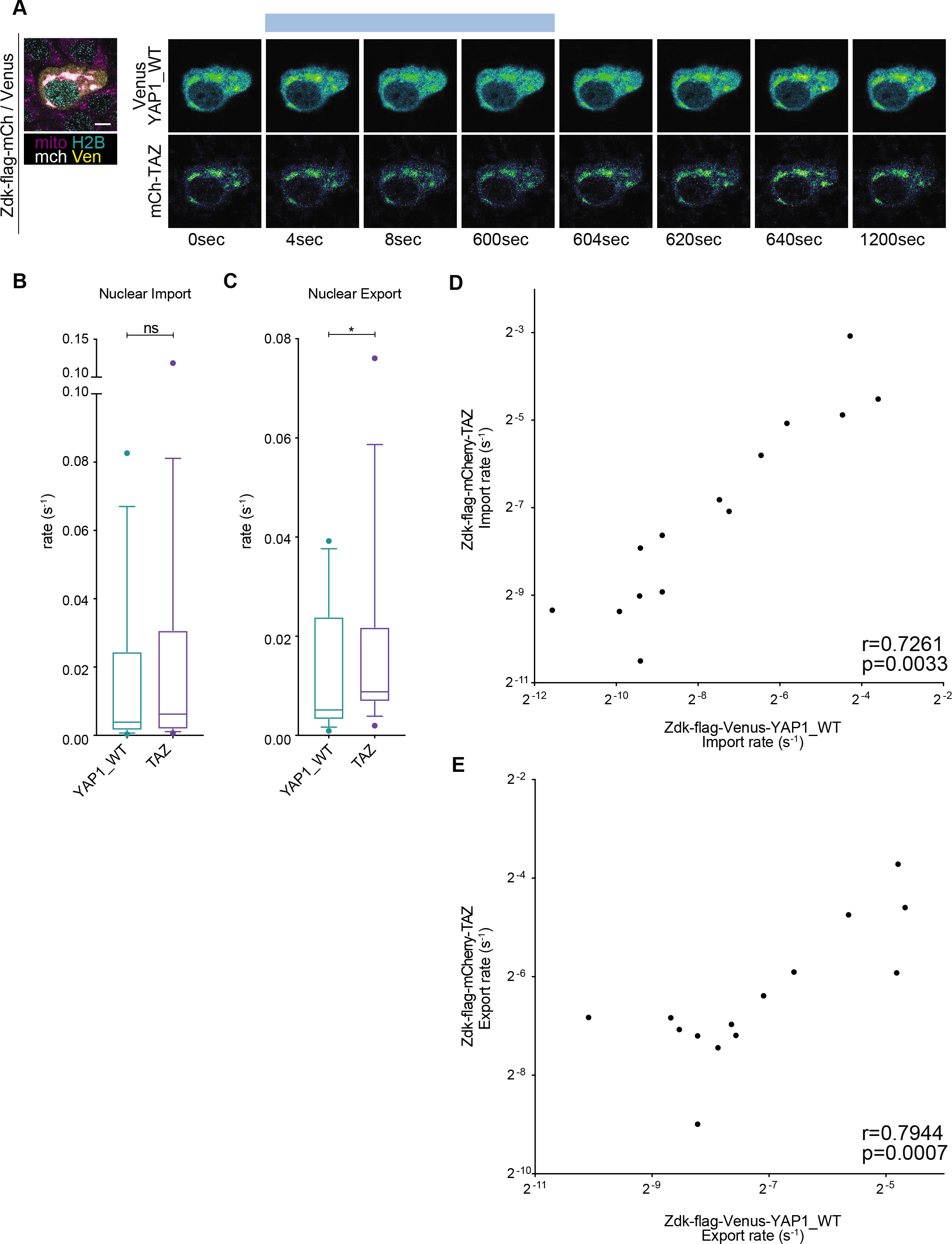
Rates of YAP1 and TAZ in the same cell. **(A)** Representative images of release and recovery experiment in Zdk-flag-Venus-YAP1_WT and Zdk-flag-mCh-TAZ using a two-colour optogenetics setup. Scale bar is 10 μm. **(B)(C)** YAP1 and TAZ nuclear import and export rates in the same cell. Boxplot (10&90 percentile, median) represents at least three independent biological replicates. Wilcoxon matched-pairs signed rank test. **(D)(E)** Scatterplot of YAP1 versus TAZ import (D) or export (E) rates in the same cell. Pearson coefficient (r) and significance (p-value) are also shown.

### Implementation of opto-release methodology on lattice lightsheet platform

YAP1 and TAZ can be associated with a wide array of binding partners in diverse sub-cellular compartments. Confocal analysis of a single optical section is not suited analyzing the distribution of molecules throughout the cell. Therefore, we sought to implement our opto-release method on a lattice lightsheet microscope with suitable laser lines for the release of the Zdk peptide from the LOV domain. The experimental design was adapted from the simple release and re-binding protocol, with additional single frames of blue light included at 80 second intervals in the re-binding phase (Fig. 6A). The additional blue light pulses were to explore if events with different kinetics might also be observed when the whole cell volume was acquired with high spatial resolution (104nm xy and 211nm z resolution). Fig. 6B shows 3D image of confluent HaCaT cells, with one expressing the Zdk-mCherry-YAP1 construct. As expected, the Zdk-mCherry-YAP1 fusion exhibits efficient localization to the mitochondrial network in frame 1, but is widely distributed throughout the cell by frame 80/480 seconds (3D view shown in Fig. 6C and xy, xz, yz orthogonal slices shown in Fig. S8A,B). Following the end of the long phase of blue light illumination, the YAP1 fusion returns to the mitochondria (Fig. 6C – 556 seconds). As expected, this localization is diminished immediately following the pulses of blue light at 560, 640, and 720 seconds (Fig. 6D left-hand panel). Quantification of the Zdk-mCherry-YAP1 signal from representative areas of the cytoplasm, and nucleus are shown in Fig. 6D. These traces clearly show the pulsatile release of the YAP1 fusion into the cytoplasm, but this does not translate into changes in nuclear signal.

**Figure 6:**
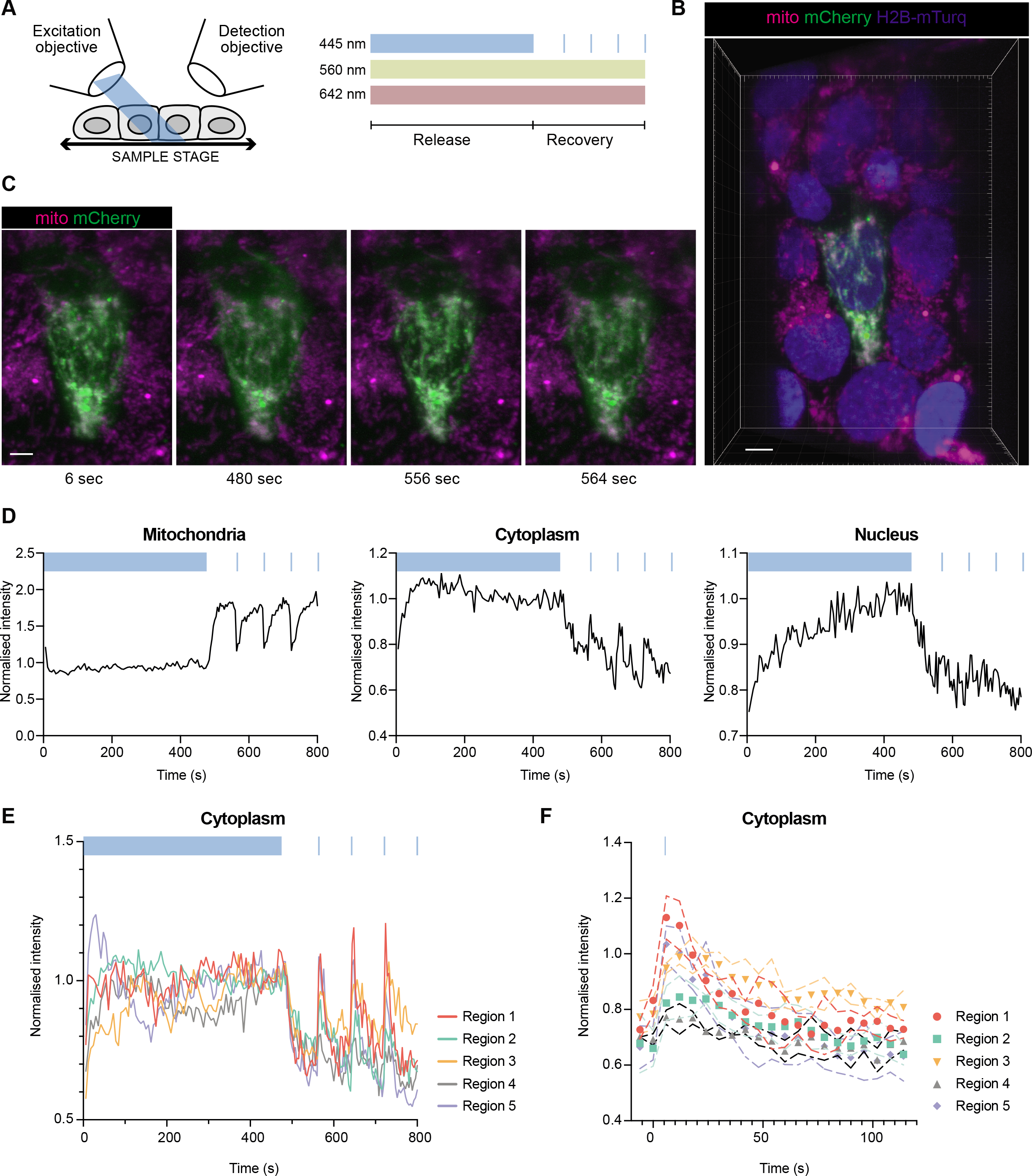
YAP1 dynamics in 3D. **(A)** Schematic representation of lattice lightsheet imaging experimental design. **(B)** Representative image of HaCat cell expressing Zdk-flag-Venus-YAP1_WT imaged on lattice lightsheet microscope. Scale bar is 10 μm. **(C)** Release and recovery experiment of Zdk-flag-Venus-YAP1_WT in 3D. Scale bar is 5 μm. **(D)** Quantification of intensities corresponding to optogenetic release and recovery experiment performed in HaCaT cell expressing Zdk-flag-Venus-YAP1_WT in 3D. The curves show the average intensities in selected regions of the mitochondria, the cytoplasm and the nucleus. **(E)** Quantification of intensities corresponding to different cytoplasmic regions during the 3D optogenetic experiment in HaCaT cell expressing Zdk-flag-Venus-YAP1_WT. **(F)** Average post-blue light spike intensity profiles for different cytoplasmic regions in HaCaT cell expressing Zdk-flag-Venus-YAP1_WT.

The purpose of performing high resolution lattice light sheet imaging was to determine if the dynamics of the YAP1 fusion protein might be varying depending on the precise subcellular localization. We therefore quantified multiple cytoplasm and nuclear regions. Interestingly, different regions showed markedly divergent kinetics. Despite the whole cell being illuminated with short pulses of blue light, the extent of gain in YAP1 signal and its duration appeared to vary from region to region. Some areas had a pronounced increase that decayed sharply (regions 1&5), while in other parts of the cytoplasm the increase was less pronounced, but more durable (regions 2&3), or barely detectable at all (Fig. 6E – similar data for the cytoplasm of a different cell is shown in Fig. S8C). Differences were also observed in the shape of the nuclear intensity profiles (Fig. S8D). To more rigorously explore the differences between cytoplasmic regions we exploited the fact that there were three equal duration pulses in the time series data. Therefore, we considered each pulse to be a ‘technical replicate’ and derived the average response for the difference cytoplasmic regions. Fig. 6F shows the difference in the average responses, with statistically significant differences in peak intensity and the rate of signal decline after the peak shown in Fig. S8E. Furthermore, a low maximal intensity post-blue light pulse correlative with a slower decay of signal; these data are consistent with slower movement of YAP1 both into and out of these regions. These data demonstrate that YAP1 dynamics vary between different regions of the cytoplasm and illustrate the utility of combined optogenetic manipulation and lattice lightsheet imaging for studying sub-cellular variation in protein dynamics.

## Discussion

Decoding the complex regulatory inputs governing the NC distribution of proteins requires accurate measurement of rates of nuclear entry and exit. During the last decade, several works showed that on YAP1 and TAZ converge an incredibly large number of different signals, from cellular architecture and microenvironment geometry to metabolic and biochemical pathways (Pocaterra et al., 2020). Here, we implement an optogenetic system to track the NC shuttling of fluorescent YAP1 and TAZ proteins coupled to mathematical modeling to derive nuclear import/export rates. This system builds upon the previously reported LOV-TRAP optogenetic tool that allows protein dissociation within a fraction of a second upon illumination with blue light (450-490 nm). In our system, which we term ‘opto-release’, we exploited the light-induced dissociation of the LOV domain tethered on mitochondria surface from a previously identified synthetic peptide (Zdk) fused to YAP1 or TAZ and then monitor their sub-cellular distribution over time. With this system we can rapidly generate a pool of fluorescently labeled cytoplasmic YAP1 and TAZ that can be tracked in the cell after release from mitochondria (light) and recovery (dark). The LOV-TRAP system has several advantages over other optogenetic systems for this purpose: i) Zdk is a small peptide and doesn’t interfere with endogenous activity of YAP1 and TAZ (as shown in Fig. S1E); ii) the LOV domain utilizes an endogenous chromophore (flavin mononucleotide), reducing external intervention during analysis; iii) it allows the study of multiple fluorescently-labeled proteins with sufficient spectral separation, such as mCherry and Venus (Fig. 5); and iv) it allows rapid, reversible and non-destructive analysis compared to FLIP, thus allowing repeated cycles of light-dark and generation of three curves, from release and recovery dynamics leading to more precise measurements.

Using our opto-release system, we measure YAP1 and TAZ shuttling rates. Despite being larger than mCherry, both YAP1 and TAZ shuttle in and out of the nucleus faster, with roughly 10% of TAZ entering or exiting the nucleus every second. This indicates active mechanisms of both import and export. The rate of YAP1 transit that we measure in epithelial HaCaT cells is similar to that which we previously reported for YAP1 in fibroblasts using FRAP and FLIP methodology (Ege et al., 2018). Direct comparison of opto-release with FLIP methods in HaCaT cells showed good concordance when the photobleaching was performed in the cytoplasm. The discordant results of nuclear photobleaching are probably due to direct photobleaching of fluorophore in the cytoplasmic regions of the cell directly above and below the nucleus. The greater height and more cuboidal nature of epithelial cells, such as HaCaT, compared to fibroblasts mean that there will be more cytoplasm above and below the nucleus. This will lead to greater levels of photobleaching occurring directly in the cytoplasm during nuclear FLIP experiments and will confound the results.

In our analysis, mutation of LATS phosphorylation target sites on YAP1 leads to reduced nuclear export that can explain YAP1_5SA nuclear accumulation (Fig. 3B), which is consistent with LATS-mediated phosphorylation being required for effective export (Ege et al., 2018). Intriguingly, the YAP1_S94A mutant, which is unable to bind TEAD, is localized in the cytoplasm at steady-state, and in our analysis its import/export rates are close to those calculated for YAP1_WT (Fig. 3A,B) even though it is more cytoplasmic than the wild-type YAP1. This could be explained if there is a pool of YAP1 that remains sequestered in the cytoplasm with binding and unbinding rates that are too slow to measure in the few minutes duration of our assays. Indeed, several studies have suggested cytoplasmic partners for YAP1 (Kanai et al., 2000, Azzolin et al., 2014, Varelas et al., 2010, Zhao et al., 2010). In the future, it will be interesting to perform longer release phase experiments with a view to the released YAP1 reaching equilibrium distribution in all sub-cellular compartments. Although we focused exclusively on NC distribution in our study, the opto-release system could equally be applied to problems involving the segregation of proteins between different cytoplasmic compartments: for example, the plasma membrane and cytoplasm. The mathematical modelling of spatial information in addition to temporal information using partial differential equations as in Ege et al. (Ege et al., 2018) will also allow us to separate import and export from motility within each compartment.

Interestingly, we could confirm that import and export rates are correlated within the same cell, but show high intercellular variability, as observed in Drosophila (Manning et al., 2018). Our conclusion that nuclear export is the main mechanism for the regulation of YAP wild-type activity differs from evidence in *Drosophila*, where Yorkie is mainly regulated by tuning nuclear import (Manning et al., 2018). A possible explanation for this discrepancy is that Yorkie (the *Drosophila* homologue of YAP1), lacks PDZ-binding motif that is required for the nuclear accumulation of YAP1 in mammals (Wang et al., 2016b, Oka and Sudol, 2009). By releasing two proteins simultaneously, we could also show that YAP1 and TAZ import and export rates co-varied indicating that they share some regulatory inputs, with LATS-mediated phosphorylation being an obvious candidate. However, we also found evidence for some specificity in regulation between YAP1 and TAZ. For example, wild-type YAP1 export rate negatively correlates with NC ratio, supporting the idea that the rate of nuclear export may be tuned in response to variation in morphological features of the cell, but this was not true for TAZ. It was also not true for YAP1_5SA, which is at least consistent with our previous finding that cytoskeletal inputs primarily tune the export rate of phosphorylated YAP1. Thus, while YAP1 and TAZ share to some extent mechanisms of NC shuttling, specific regulations are in place, as shown by a recent study on TAZ-specific NES/NLS sequences (Kofler et al., 2018). Of note, FRAP early experiments with GFP-TAZ showed that TEAD-binding defective TAZ mutants exhibit higher nuclear export (Chan et al., 2009). Despite correlation of import/export with several morphological metrics, we couldn’t observe a link between NC shuttling and nuclear deformation, as reported in Elosegui-Artola et al. (Elosegui-Artola et al., 2017). However, we did not apply any direct or indirect perturbations to the nuclear envelope.

Current FRAP and FLIP methodologies are routinely employed with a focus on a single optical plane of the cell. This is partly due to the difficulties in obtaining multiple optical sections with the necessary speed and partly due to the difficulties accounting for the changes in the size and intensity of the photobleaching region as a function of distance in the z-dimension. However, opto-release technology does not suffer from the variation in photobleaching as a function of z position, especially if the whole cell is illuminated. Furthermore, advances in lattice lightsheet technology now allow for the rapid acquisition of high resolution volumetric data of whole cells. Therefore, we implemented our opto-release methodology on a lattice lightsheet platform. This revealed additionally complexity in the behavior of the Zdk-mCherry-YAP1 fusion in different regions of the cytoplasm, with some regions showing rapid increases and decreases in signal following a pulse of blue light and other regions showing lower peak intensities and slower decreases in signal. The regions with slower kinetics may reflect areas with slow diffusion or slow binding to a tethered partner molecule. In the future, it will be interesting to characterize these different regions by co-staining for other organelles and YAP1 binding partners. These differences also highlight the limitations of applying a simple mathematical model that only considers mitochondrial, cytoplasmic, and nuclear compartments with no locally varying properties or sub-compartments. This is likely a source of uncertainty or error in our fitting of import and export rates. Nonetheless, ODE modelling of the predicted YAP1 levels in different cellular compartments under the blue light regimen used in the lightsheet experiments was consistent with the observed changes in mitochondrial and nuclear Zdk-mCherry-YAP1 signal, and with cytoplasm regions 1 and 5 (compare Fig. S8F and Fig. 6E). The modelling also predicts that short pulses of blue-light would have only a minimal effect on nuclear intensity. The smaller magnitude of changes in the nucleus mean that the experimental data are noisier, nonetheless, they support the prediction of minimal effect of the blue-light pulses on nuclear YAP1 levels. Solving the issue of heterogenous YAP1 dynamics in the cytoplasm will require a more complex model with a larger number of compartments and more spatial features. It will also require implementation of 3D segmentation tools for different cellular compartments.

In the present study, we report the use of a reversible optogenetic system based upon the LOV-TRAP system (Wang and Hahn, 2016, Wang et al., 2016a). This methodology allows controlled release of fluorescent proteins that can be tracked over time in the different cell compartments. The reversibility of the optical release enables more complex experimental design than is possible with conventional FRAP and FLIP analyses. In particular, by varying the timing and length of release and re-binding phases it is possible to acquire data about processes happening with different timescales in the same experiment. Alternatively, repetitive use of a signal illumination pulse or sequence can effectively generate ‘technical replicates’ in a single experiment, leading to more accurate measurements. We generated a MATLAB app for thresholding of different cellular compartments and compensation of cell motion. This tool then extracts numerical values for NC localization that are used by differential equations to generate import/export rates. While we have used this system to gain further insights in YAP1 and TAZ NC shuttling, the modular nature of the constructs will make it easy to adapt to a range of transcription factors and regulators, including SMADs, NF-kB, MRTF, IRFs, and STATs. We anticipate that opto-release systems based on this framework will facilitate the understanding of how diverse inputs regulate the sub-cellular localization of proteins and will be of wide utility to cell biologists.

---

**Supplementary Figure 1:**
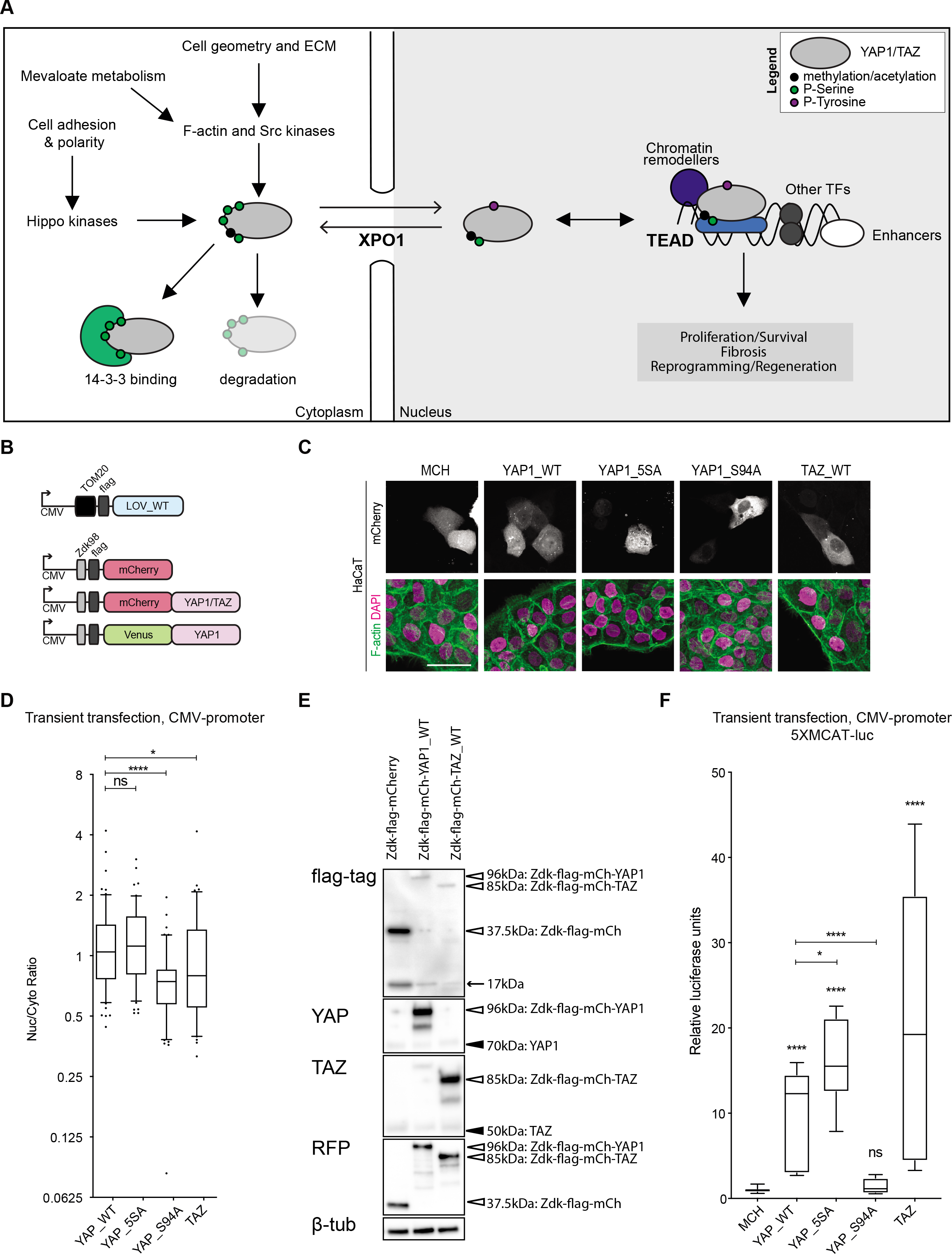
Related to Figure 1. Localization, expression and function of unsequestered Opto-release constructs. **(A)** Schematic of main known mechanisms of regulation of YAP1 and TAZ. **(B)** Schematic of various constructs used for opto-release system. **(C)** Representative images of HaCaT cells expressing CMV-driven Zdk constructs (transient transfection). Actin is presented in green, DAPI in magenta and mCherry in greyscale. Scale bar is 50 μm. **(D)** Boxplot (10&90, median) of nuclear-to-cytoplasmic ratio (log2 scale) corresponding to three experimental repeats of HaCaT cells transiently transfected with CMV-driven Zdk constructs (n>29cells; n>20 for Taz, 2 experimental repeats). Mann-Whitney U test. **(E)** Expression of unsequestered Zdk constructs. Western blot showing expression of Opto-release constructs stably expressed in HaCaT cell line. Empty arrowheads point to Opto-release protein (Zdk-flag-mCh 37.5kDa, Zdk-flag-mCh-YAP1_WT 96kDa, Zdk-flag-mCh-TAZ_WT 85kDa) and filled arrowheads point to endogenous protein (YAP1 70kDa, TAZ 50kDa). An arrow points to a cleavage product. Anti-RFP antibody recognizes mCherry fluorescent protein. [Symbol]-tubulin as loading control (42kDa) is also presented. **(F)** Luciferase assay using TEAD-driven reporter (5xMCAT) for YAP1/TAZ activity in HaCaT cells following transient transfection with Zdk constructs driven by CMV promoters. Boxplot (10&90, median) represents three independent experiments, each with 3 technical replicates. Mann-Whitney U test, statistics above each bar represent comparison to MCH negative control.

**Supplementary Figure 2:**
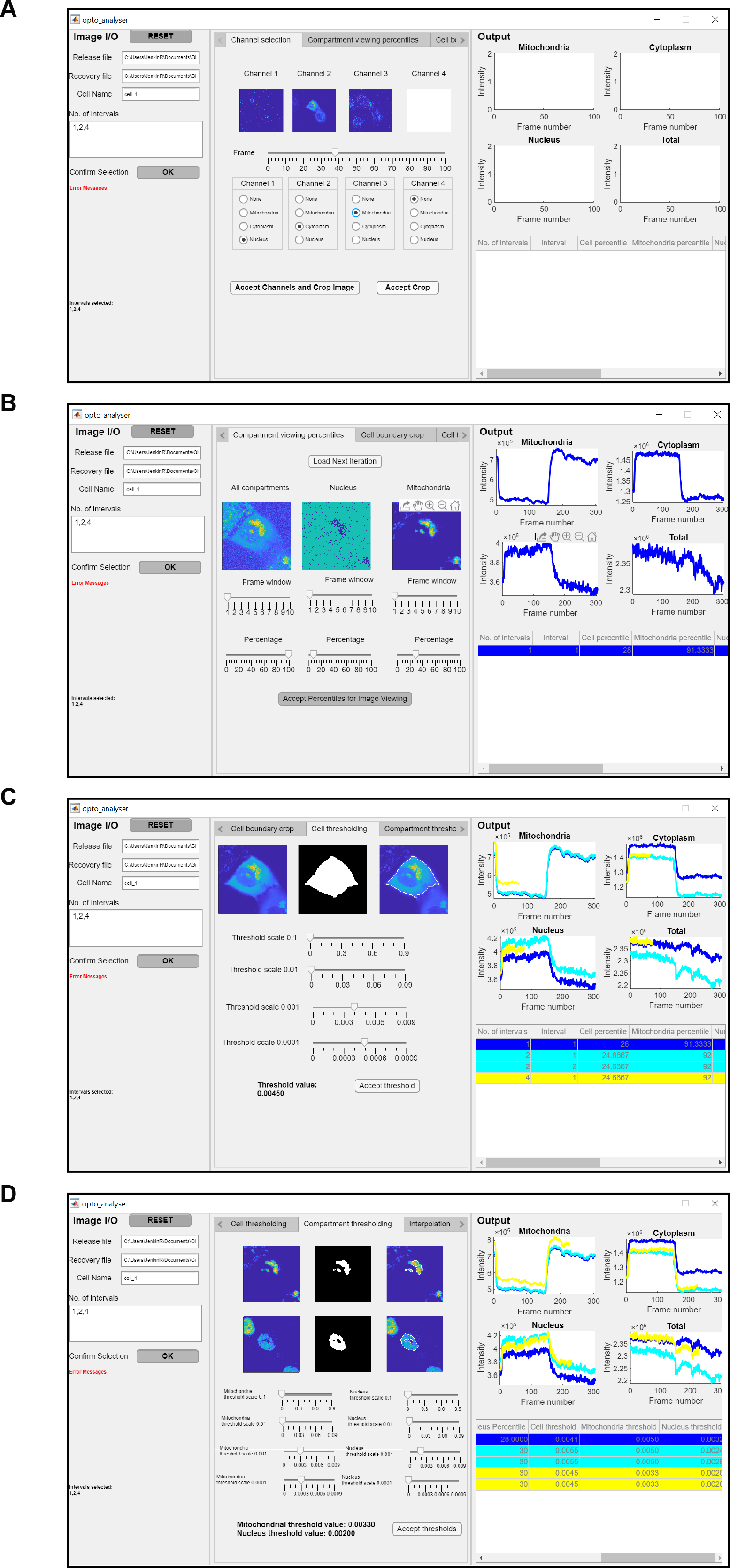
Related to Figure 1. Image analysis pipelines. **(A)** MATLAB app showing user input and channel selection tab. **(B)** MATLAB app showing user determined percentile projections for each channel in the central panel. Here, the percentile for the whole cell is too high, leading to high noise and interference from neighbouring cells. The nuclear channel’s percentile is too low, leading to no obvious nuclear object. The percentile for the mitochondrial channel is close to optimum. The right-hand panel illustrates population intensity profiles and segmentation information for a single window split. **(C)** MATLAB app showing, in the central panel, how the threshold value to segment the whole cell can be altered. The right-hand panel shows population intensity output and relevant parameter values for the movie split into a single window (blue), two windows (cyan) and the first window of a four window split (yellow). **(D)** MATLAB app showing the segmentation of the nucleus and mitochondria in the central panel and further population intensity output and relevant parameter values for one (blue), two (cyan) and four (yellow) window splits in the right-hand panel. Changes in thresholding in the central panel will change the intensity profiles of the third quarter of the yellow signal (four window split) in real time. The intensity profile of the nucleus is higher in this quarter than the one preceding it, suggesting that the nuclear threshold should be higher.

**Supplementary Figure 3:**
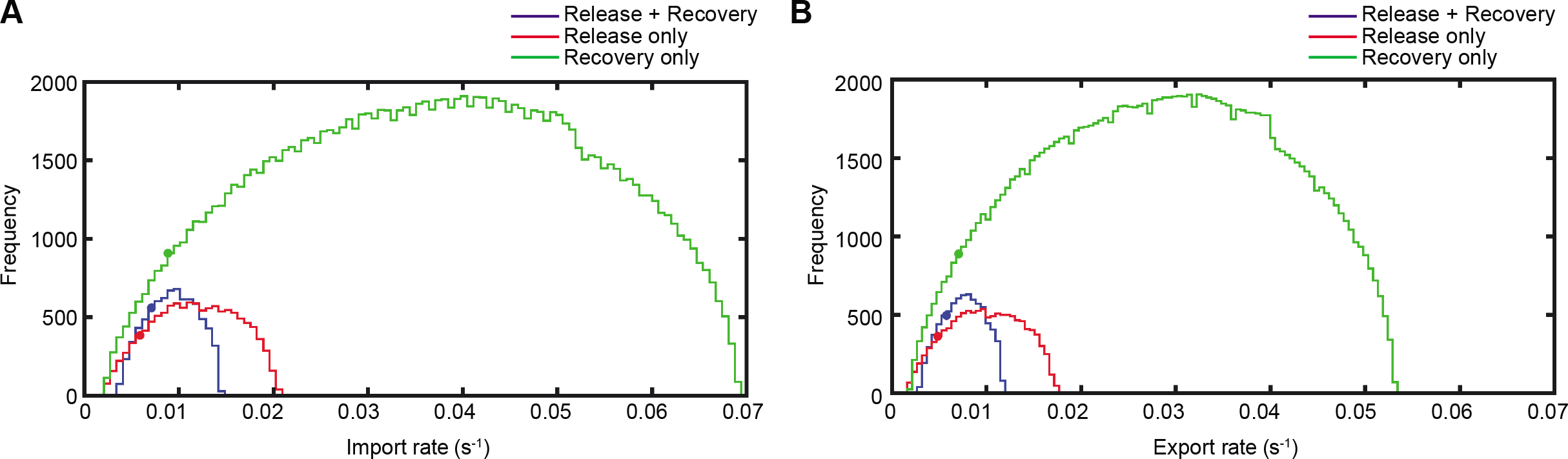
Related to Figure 2. Analysis of uncertainty of parameter values for release fit, recovery fit or release + recovery fit. **(A)(B)** Frequency distributions of import (A) and export (B) values that produce fits with sum of squares of error at most twice that of the global optimum when varying import and export simultaneously. The dots show the location of optimum fit and suggest release-only fitting underestimates both import and export while recovery-only overestimates import and export. All other parameter values were fixed at their global optimums. Size of distribution shows greatest certainty with release and recovery and least with only recovery.

**Supplementary Figure 4:**
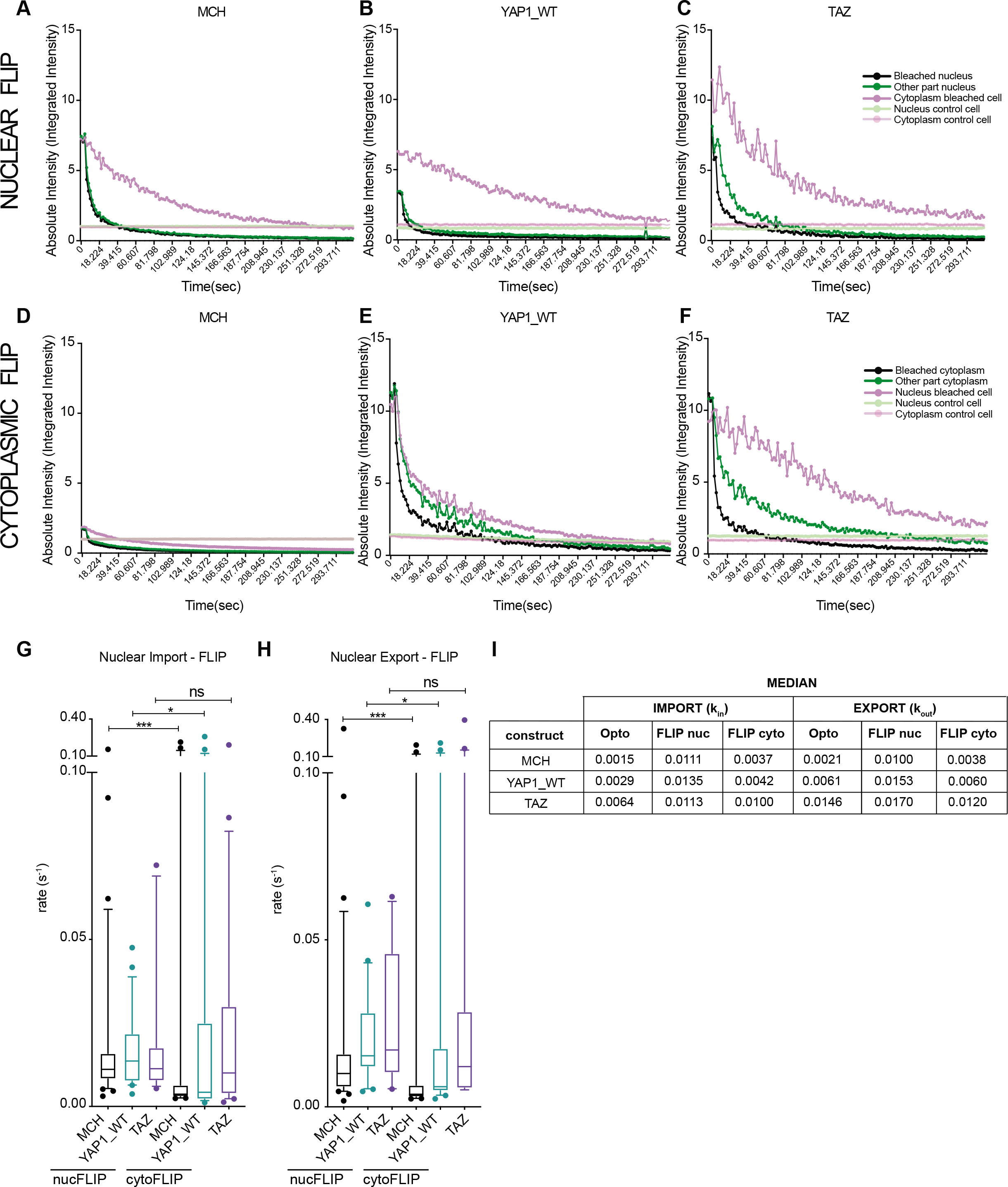
Related to Figure 2. Confirmation of shuttling rates using FLIP. **(A)(B)(C)** Nuclear FLIP analysis of unsequestered Zdk-mCherry constructs. Quantification of absolute intensities normalized to control cell and background intensity, corresponding to FLIP experiment performed in the nucleus of HaCaT cells transiently transfected with (A) Zdk-flag-mCherry (n=32 cells, 4 biological replicates), (B) Zdk-flag-mCh-TAZ (n=17 cells, 3 biological replicates), (C) Zdk-flag-mCh-YAP1_WT (n=21 cells, 3 biological replicates). Only the mean of all cells is shown for clarity. **(D)(E)(F)** Cytoplasmic FLIP analysis of unsequestered Zdk-mCherry constructs. Quantification of absolute intensities normalized to control cell and background intensity, corresponding to FLIP experiment performed in the cytoplasm of HaCaT cells transiently transfected with (D) Zdk-flag-mCherry (n=22 cells, 3 biological replicates), (E) Zdk-flag-mCh-TAZ (n=22 cells, 3 biological replicates), (F) Zdk-flag-mCh-YAP1_WT (n=20 cells, 3 biological replicates). Only the mean of all cells is showed for clarity. **(G)(H)** YAP1 and TAZ nuclear import and export rates acquired using FLIP methods. The numerical values of nuclear import (*k_imp_*) and export (*k_exp_*) rates for Zdk-flag-mCherry, Zdk-flag-mCh-TAZ, Zdk-flag-mCh-YAP1_WT are shown. Boxplot (10&90 percentile, median) represents at least 3 independent biological replicates. Mann-Whitney U test. **(I)** Table comparing medians of shuttling rates derived using Opto-release method, FLIP nucleus and FLIP cytoplasm.

**Supplementary Figure 5:**
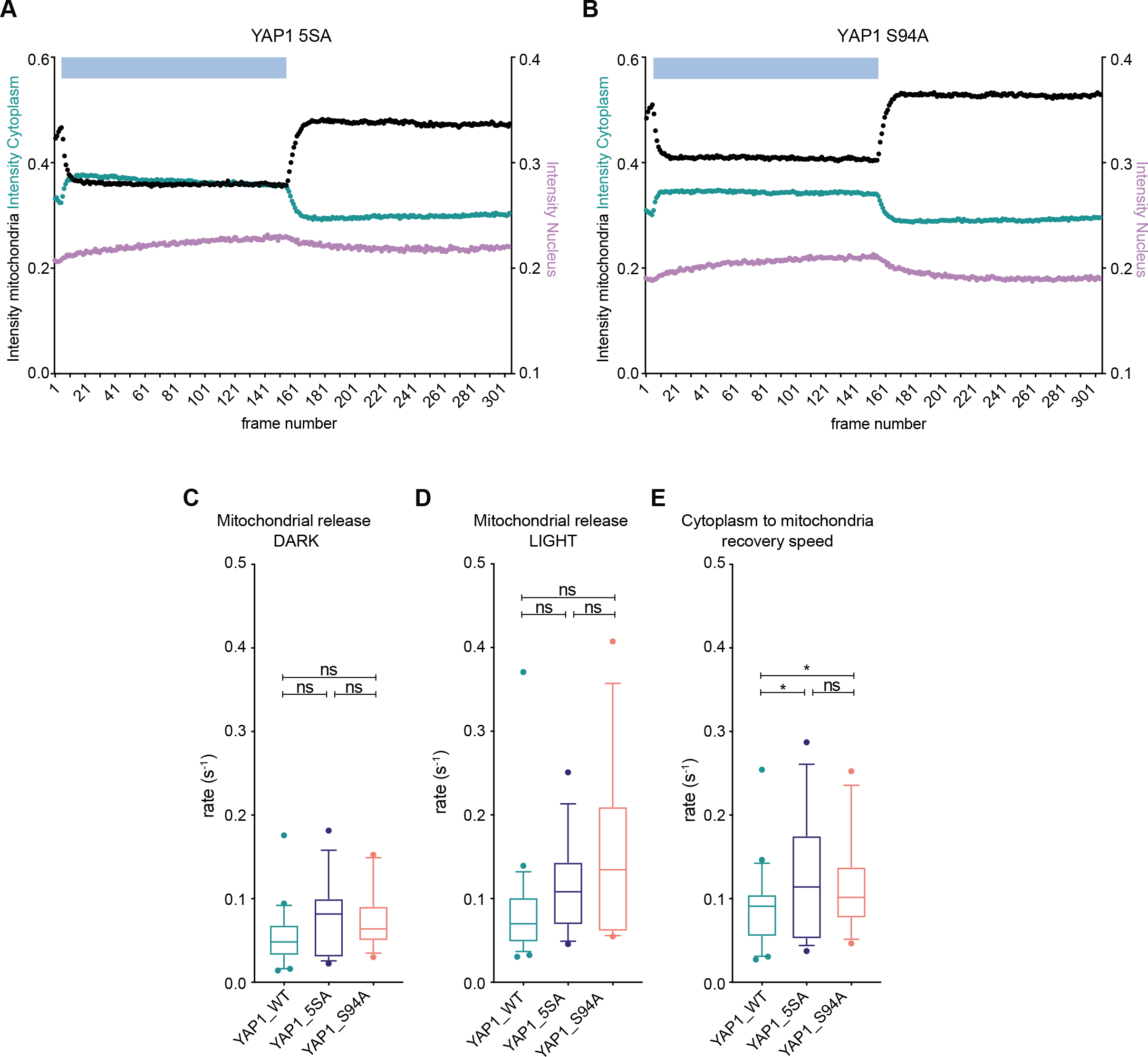
Related to Figure 3. YAP1 mutants parameters. **(A)(B)** Quantification of absolute intensities corresponding to optogenetic release and recovery experiment performed in HaCaT cells transiently transfected with (A) Zdk-flag-mCh-YAP1_5SA (n=24) or (B) Zdk-flag-mCh-YAP1_S94A (n=22). Only the mean of all cells from at least 3 biological replicates is shown for clarity. Intensity has been corrected for bleaching and background signal. **(C)(D)(E)** The numerical values of mitochondrial sequestration rates for Zdk-flag-mCherry-YAP1_5SA (n=14) and Zdk-flag-mCherry-YAP1_S94A (n=12) compared to Zdk-flag-mCherry-YAP1_WT reproduced from Figure 2 (n=28), in steady state (darkness), during release and during recovery phases. Number of cells per each condition is given in brackets. Boxplot (10&90 percentile, median) represents at least three independent biological replicates. Mann-Whitney U test.

**Supplementary Figure 6:**
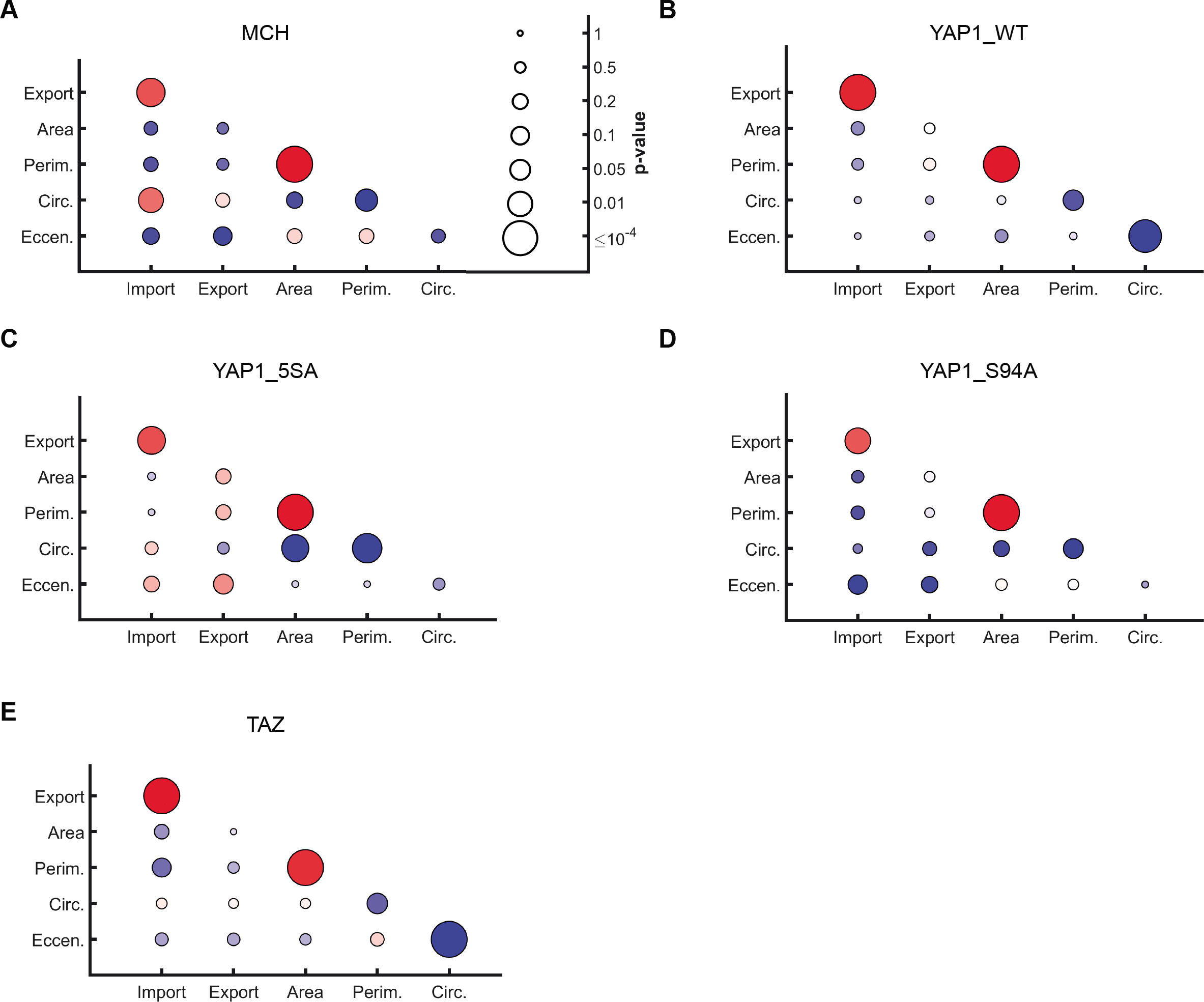
Related to Figure 4. Cytoplasm morphology correlation with import and export. **(A) (B)(C)(D)(E)** Import and export correlations with cytoplasmic morphology, specified by area, perimeter, circularity and eccentricity. Plots show Zdk-flag-mCherry (A), Zdk-flag-mCh-YAP1_WT (B), Zdk-flag-mCh-YAP1_5SA (C), Zdk-flag-mCh-YAP1_S94A (D) and Zdk-flag-mCh-TAZ (E). Circle colour reflects Pearson correlation (bright red +1, dark blue −1) and circle size the p-value of the correlation (large, significant; small, nonsignificant).

**Supplementary Figure 7:**
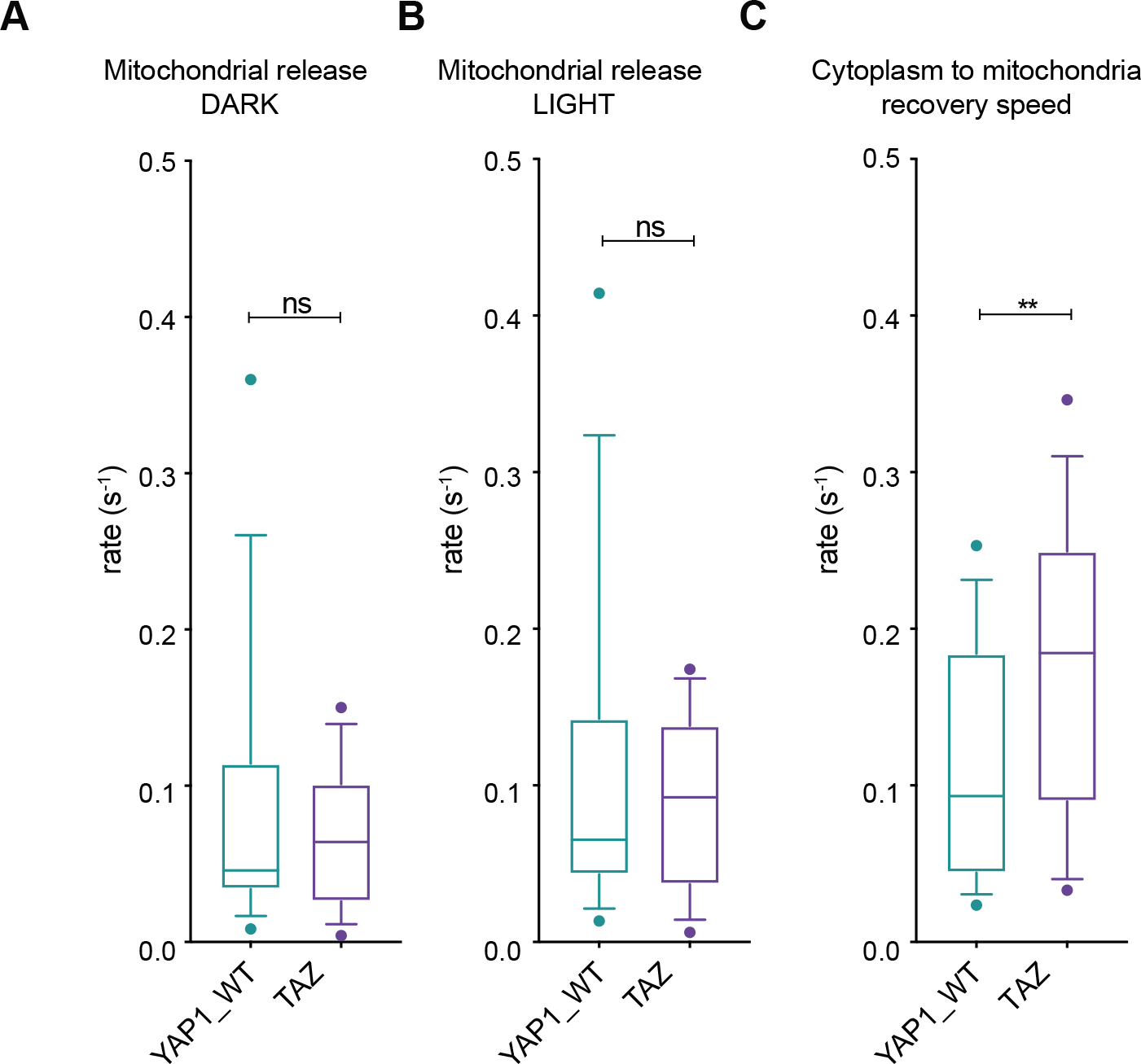
Related to Figure 5. Rates of YAP1 and TAZ in the same cell. **(A)(B)(C)** The numerical values of mitochondrial sequestration rates for Zdk-flag-Venus-YAP1_WT and Zdk-flag-mCherry-TAZ (n=14 cells) compared in the same cell in steady state (darkness), during release and during recovery phases. Number of cells per each condition is given in brackets. Boxplot (10&90 percentile, median) represents at least three independent biological replicates. Wilcoxon matched-pairs signed rank test.

**Supplementary Figure 8:**
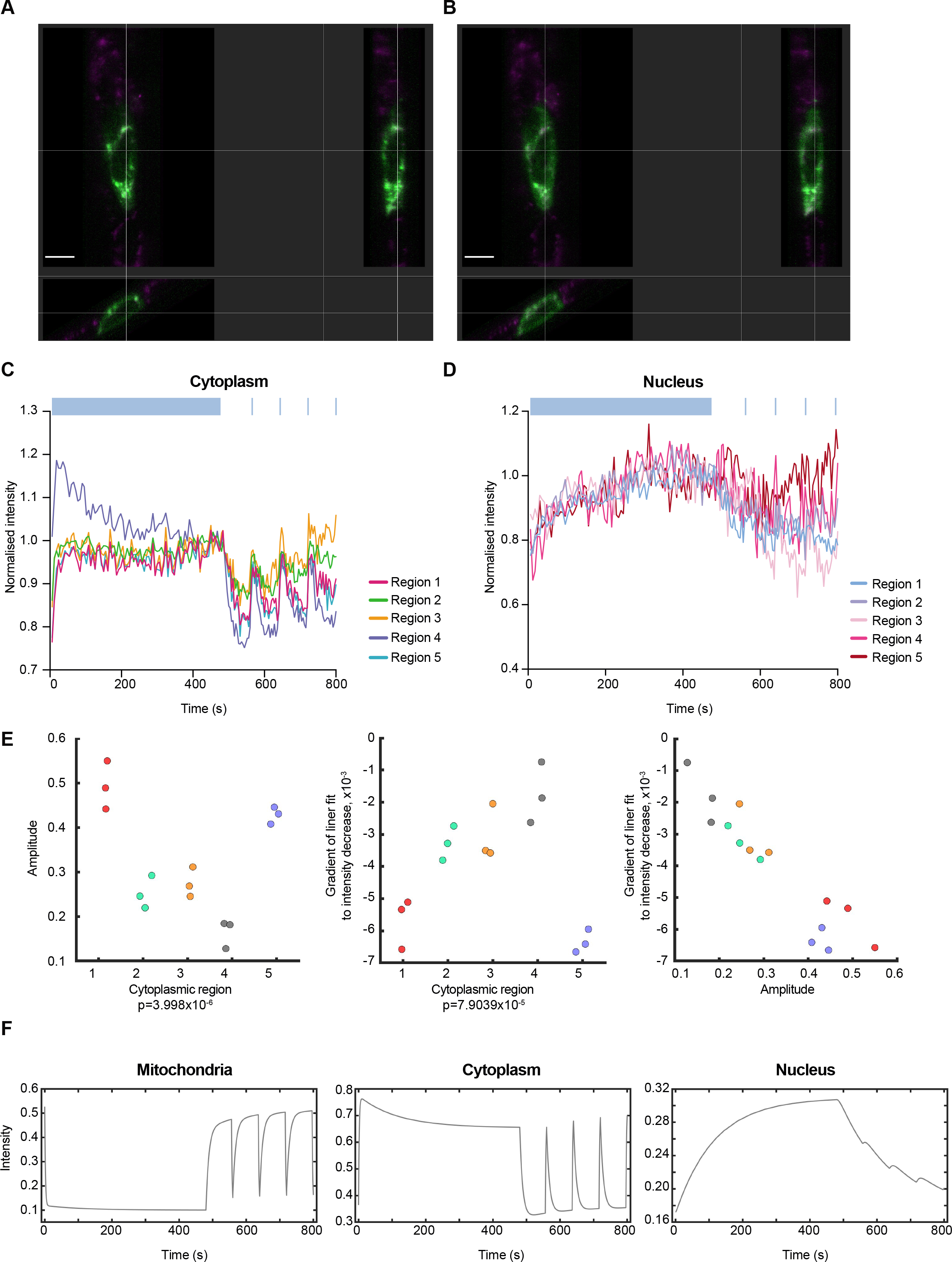
Related to Figure 6. YAP1 dynamics in 3D. **(A)(B)** Orthogonal views of HaCaT cell expressing Zdk-flag-Venus-YAP1_WT fusion during recovery (A) and release (B). Scale bar is 10 μm. **(C)** Quantification of intensities corresponding to different nuclear regions during the 3D optogenetic experiment in HaCaT cell expressing Zdk-flag-Venus-YAP1_WT. **(D)** Quantification of intensities corresponding to different cytoplasmic regions during the 3D optogenetic experiment in another HaCaT cell expressing Zdk-flag-Venus-YAP1_WT (additional to Fig. 6E). **(E)** Scatter plots of peak amplitude and gradient of decrease from maximum value of each peak for each cytoplasmic region of data in Fig. 6E (left and centre) and scatter plot of amplitude of peaks versus gradient of decrease (right). Peak amplitude approximated by maximum minus minimum signal value. For the gradient of decrease a linear fit was made to all data points after and including the peak value. The gradient provides an approximation to rate of decrease of the peak. Data shows region dependent amplitudes, persistence of peaks and that peak amplitude and persistence of peak are correlated. Statistical tests carried out using one-way ANOVA. **(F)** Using the model of Fig 1D to predict functional behaviour of population intensity in the mitochondria, cytoplasm and nucleus for the lightsheet experimental set-up. Parameter values are those fitted to YAP1_WT. Release rate during illumination has been increased by a factor of five to qualitatively match the increase in bleaching rate in the lightsheet data. The predictions show good qualitative agreement with Fig 6D.

## Supplementary Movies

Movie 1: Movie shows the first 60 frames (240 seconds) of the blue light illuminated release phase of Zdk-mCherry and the first 60 frames (240 seconds) of the non-illuminated recovery phase.

Movie 2: Movie shows the first 50 frames (200 seconds) of the blue light illuminated release phase of Zdk-YAP1-mCherry and the first 50 frames (200 seconds) of the non-illuminated recovery phase.

Movie 3: Movie shows the first 50 frames (200 seconds) of the blue light illuminated release phase of Zdk-TAZ-mCherry and the first 50 frames (200 seconds) of the non-illuminated recovery phase.

Movie 4: Movie shows 3D image of a HaCaT cell release and recovery experiment on the lattice lightsheet microscope.

## Materials and methods

### Cell lines and cell culture

HaCaT immortal keratinocyte cells were acquired from the Cell Services Facility at the Francis Crick Institute. H2B-mTurquoise2 was introduced to cells using the PiggyBac transposon system, where the plasmid of interest is transfected together with a transposase plasmid PBase, and was described before (Ege et al., 2018). Cells expressing the construct were selected using Puromycin. All the other plasmids were introduced transiently. Cells were cultured in DMEM (Gibco) with 10% foetal bovine serum (FBS) (PAA Labs) and 1% penicillin-streptavidin (Pen-Strep, Gibco) at 37°C and 5% CO2. The cells were removed from the cell culture dish with 0.1% trypsin and 0.02% versene (both Life Technologies) after washing with PBS (Phosphate buffered saline, 1.5 mM KH2PO4, 8 mM Na2HPO4, 2.7 mM KCl, 137 mM NaCl, pH 7.4). Trypsin was neutralised by addition of cell culture media containing 10% FBS.

### Plasmids

All the plasmids generated for the opto-release system were cloned using the Gibson Assembly System (NEB) by combining pRK5.1 backbone with desired PCR fragments according to the manufacturer’s instructions. It is based on a previously published LOV-TRAP (Wang and Hahn, 2016, Wang et al., 2016a). The photo-switchable modified common oat (Avena sativa) light-oxygen-voltage (LOV) 2 domain from Phototropin 1 protein is fused to TOM20 protein fragment, which embeds it in the outer mitochondrial membrane. On the other side of the system there is the YAP1/TAZ protein, which is joined to Zdk, an engineered small protein domain based on the Z domain described previously (Wang and Hahn, 2016, Wang et al., 2016a), and fluorescent proteins (mCherry or Venus, both photostable in this system and in the timescale of our experiments. YAP1 corresponds to the human YAP1-2γ isoform, which is 504 amino acids long (Sudol, 2013) and the mutants were described before (Ege et al., 2018). The five serines mutated for the 5SA mutant correspond to serines 61, 109, 127, 164 and 397 (=381) for the isoform used in this study. The two plasmids used for the luciferase experiments, pGL3-49 and pGL3-5xMCAT-49, a gift from Nic Tapon. The pPB-puro-H2B-mTurquoise2 was generated in the lab (Ege et al., 2018).

| Plasmid name | Colours | Promoter | Use | Source |
| --- | --- | --- | --- | --- |
| pRK5.1-TOM20-flag-LOVwt | none | CMV | optogenetics | cloned |
| pRK5.1-Zdk-flag-mCh | mCherry | CMV | optogenetics | cloned |
| pRK5.1-Zdk-flag-mCh-YAP1_WT | mCherry | CMV | optogenetics | cloned |
| pRK5.1-Zdk-flag-mCh-YAP1_5SA | mCherry | CMV | optogenetics | cloned |
| pRK5.1-Zdk-flag-mCh-YAP1_S94A | mCherry | CMV | optogenetics | cloned |
| pRK5.1-Zdk-flag-mCh-TAZ | mCherry | CMV | optogenetics | cloned |
| pRK5.1-Zdk-flag-Venus-YAP1_WT | mCherry | CMV | optogenetics | cloned |
| pGL3-49 | luciferase | N/A | luciferase assay | Tapon lab |
| pGL3-5xMCAT-49 | luciferase | N/A | luciferase assay | Tapon lab |
| pcDNA3-Clover | Clover | CMV | filler plasmid for luciferase assay | Addgene #40259 |
| pGL4.75 CMV-Renilla | renilla | N/A | luciferase assay | Promega, #E6931 |

### DNA Transfection

For transient transfection with plasmid DNA, cells were transfected with Lipofectamine LTX and Plus reagents combination (# 15338100, Thermofisher) according to the manufacturer’s instructions. One day prior to the transfection, cells were seeded 0.5-1x 10^6^ cells for Western blotting and luciferase transfection in a 6 well plate, or 20 000-100 000 for live imaging and immunofluorescence in a 24 well plate. Prior to transfection media was changed to Pen-Strep free. On the day of transfection two tubes were prepared (proportions below for 24 well plate):

Tube 1: 100 μl OptiMEM + 1-2 μg DNA + 2 μl Plus reagent
Tube 2: 100 μl OptiMEM + 4 μl Lipofectamine LTX

The two tubes were mixed and incubated for 5 minutes separately, and then 5 minutes together. The transfection mix was added drop wise to cells and incubated for 4-6 hours at 37 °C. Subsequently, the transfection mix was removed and media with Pen-Strep was added. Performing transfection without Pen-Strep is very important and greatly affects transfection efficiency. If the cells are kept longer than 6 hours in the transfection mix, cell viability can become compromised.

### Immunofluorescence

All immunofluorescence experiments were performed on cells seeded on glass in MatTek plates (MatTek Co., Ashland, MA, USA). Cells were fixed in 4% paraformaldehyde, washed with 0.01% Triton X100 PBS and permeabilised by incubation in 0.2 % Triton X100, PBS for 20 minutes at room temperature. Samples were subsequently blocked for 1 hour with 5%BSA, 0.01% Triton X100 PBS before incubation with Phalloidin-Atto633 (Sigma, # 68825-10NMOL) and DAPI (Sigma, # d9542-5mg) in 5%BSA, 0.01% Triton X100 PBS 1 hr at RT. Primary antibody was washed off in 3 washes of 5 minutes with 0.01% Triton X100, PBS, and then retained in PBS until imaging.

### Luciferase assay

Luciferase assays were performed with the dual luciferase assay kit (Promega). Cells were lysed using passive lysis buffer. Lysates were placed into a white 96-well plate (Perkin Elmer) to assess Luciferase and Renilla activities using Envision Multilabel plate reader (Perkin Elmer). To normalize, the measurements of firefly luciferase activities were normalized to the renillla luciferase activities of the same sample.

### Western Blot

For protein analysis, cells were lysed with 1 x SDS sample buffer (0.32 M Tris pH 6.8, 10% SDS, 50% glycerol, 3M β-mercaptoethanol, 0.05% bromophenol blue) added directly to wells, and then cells were scraped and collected. Each sample was sonicated and then boiled at 95 °C for 5min before being used for Western blotting. Samples were separated on SDS-PAGE gels (Bio-rad). A pre-stained Dual Colour protein ladder (Bio-rad, #1610374) was run with the samples. The proteins in the gel were transferred to polyvinylidene fluoride (PVDF) membrane (GE Healthcare Life Sciences). The membranes were blocked with 2% milk (Marvel) or 2%BSA in TBST (137 mM NaCl, 2.7 mM KCl, 20 mM Tris HCl, pH 7.4, 0.01% Tween 20) for 1 hour at RT, and then incubated with primary antibody at 4°C overnight in 0.1% milk/BSA TBST. After 3 washes with TBST, membranes were incubated 45 min at RT with HRP-labelled secondary antibody in 0.5% milk TBST (1:50 000, Fisher Scientific). The blot was then developed by rinsing it for 1 minute in Luminata WesternHRP Substrate (Millipore) before being exposed in the ImageQuant600RGB machine (GE Healthcare Life Sciences). The list of antibodies used for Western blotting is outlined below.

| Antibody | Species | Source | WB | IF |
| --- | --- | --- | --- | --- |
| YAP1 (D8H1X) | rabbit mono IgG | Cell Signalling Technology, #14074 | 1:1000 | 1:500 |
| TAZ (V386) | rabbit poly | Cell Signalling Technology, #4883 | 1:1000 | 1:200 |
| β-tubulin | mouse mono IgG1 | Sigma, #T7816 | 1:5000 | 1:1000 |
| RFP (5F8) | rat mono IgG2a | Chromotek, #5F8-100 | 1:1000 | N/A |
| flag tag | mouse mono IgG1 | Sigma, #F1804 | 1:1000 | N/A |

### Statistical analysis

Statistical analyses were performed using Prism software (GraphPad Software). p-values were obtained using Mann-Whitney unpaired t–test or Wilcoxon matched-pairs signed rank test, with significance set at p < 0.05. Graphs show symbols describing p-values: *, p < 0.05; **, p < 0.01; ***, p < 0.001, ****, p<0.0001; ns, not significant.

### FLIP imaging

Cells were plated at low confluence and cultured overnight in glass bottom MatTek dishes. Cells were transfected, and 24-48 hrs after DNA transfection imaged. One hour prior to imaging, the medium was changed to Leibovitz L-15 media with 10% serum. The cells were subsequently imaged with a Zeiss LSM880 microscope equipped with an argon laser (Zeiss, Germany) and a 63X objective (Zeiss, alpha-Plan Apochromat 63x/1.46 NA oil korr TIRF). For FLIP experiments, bleaching was performed using the laser at 100% capacity with a wavelength corresponding to fluorophore excitation wavelength at a single square ROI of 8×8 pixels (4.4 mm2). All images were 12-bit and 256×256 pixels. Before photobleaching, 3 measurements of fluorescence were taken. The ROI was then photobleached between every frame for 2 seconds using maximum laser power. A series of 150 images were taken every 2 seconds for up to 5 minutes. For nucFLIP, bleaching region was placed in the nucleus, whilst in the cytoplasm for cytoFLIP.

### Optogenetics light activation

Cells were plated at low confluence and cultured overnight in glass bottom MatTek dishes. Cells were transfected, and 24-48 hrs after DNA transfection imaged. One hour prior to imaging, the media was changed to Leibovitz L-15 medium with 10% serum supplemented with MitoTracker Deep Red (10 000x, #M22426) for staining the mitochondria. Half an hour prior to imaging, the medium was changed again to Leibovitz L-15 medium with 10% serum without the stain.

The cells were subsequently imaged with a Zeiss LSM880 microscope equipped with an argon laser (Zeiss, Germany) and a 40X objective (Zeiss, Plan Apochromat 40x/1.3 NA oil korr DIC M27) with an environmental chamber set at 37°C. Optogenetic activation was performed using 458 nm laser line at 1% laser power on the bleaching function with 50 iterations selecting an area encompassing the whole cell. Before activation, 5 frames were imaged without photoactivation/bleaching function. Then, 150 frames of release with photoactivation and 150 frames of recovery without photoactivation were acquired. Frames were acquired approximately 4 sec apart, as this corresponds to the approximate speed of light recovery of AsLOV2 domain (Strickland et al., 2012, Strickland et al., 2010). In order to simultaneously acquire two or three other channels, a beam splitter 458/514/561/633 was used, which allows imaging in the cyan, yellow, red and far-red colours compatible with light activation at 458 nm. Nuclei are imaged with very low 458 nm laser power which did not interfere with light activation at the same wavelength. Mitochondria are imaged in the far-red spectrum. The proteins of interest can be placed in the yellow (Venus) or the red (mCherry) channels.

### 3D optogenetics light activation

All the experiments in 3D were performed on LLSM (lattice light sheet microscope) in the Advanced Imaging Center at the Howard Hughes Medical Institute, Janelia Research Campus. HaCaT cells were seeded on uncoated coverslips (CS-5R; Warner Instruments) in a 30 ul droplet 10 000-30 000 cells/droplet and left to settle for 1hr before filling the plate well with media. Cells were transfected as in 2D condition.

The LLSM system was configured and operated as previously described (Chen et al., 2014). Samples were kept at 37°C and illuminated using a 445 nm laser (Oxxius diode laser, initial laser power 132 mW) at 100% acousto-optic tunable filter transmittance, 560 nm laser (MPB fiber laser, rated 500 mW) at 100% acousto-optic tunable filter transmittance and 642 nm laser (MPB fiber laser, rated 500 mW) at 10% acousto-optic tunable filter transmittance. Excitation objective was Special Optics 0.65 NA lens and detection objective was Nikon CFI Apo LWD 25x–Water dipping, 63x magnification, 1.1 NA, signal detected on two Hamamatsu Orca Flash 4.0 v2 sCMOS cameras. To separate the signal between two cameras, for channel detecting MitoTracker DeepRed we used Camera A BLPO-647R-25 long band pass filter. For Camera B we used NF03-t42E-25 (Notch filter), BLP01-458R-25 long pass and FF01-465/537/623-25 beam splitter. Signal was split with a dichroic mirror FF640-FDi014 to detect H2B-mTurquoise and mCherry. For the release phase 80 frames with 445 nm laser on were acquired, and in the recovery phase 4 cycles of 19 frames without and 1 frame with 445 nm laser were imaged (“staggered release”). In addition providing novel information about protein dynamics, staggered release can be used to map the position of the nucleus. The software used to operate the instrument and collect data was LabView (National Instruments). FIJI was used for the quantification of fluorescent intensity in different regions of interest. FIJI was also used to convert files into .h5 format for subsequent 3D visualization and the generation of movies using Imaris.

### Immunofluorescence quantification for nuclear-to-cytoplasmic ratios

For quantification of subcellular localization of protein of interest, the nuclear-to-cytoplasmic ratio was calculated manually. For each cell, a single region of interest (ROI) square of 8×8 pixels was placed in the nucleus and cytoplasm. ROIs were always selected in the DAPI channel of IF images to avoid bias. The cytoplasmic ROI was then confirmed to be in the correct cell by checking the actin channel. The nuclear ROI was divided by the cytoplasmic ROI to derive nuclear-to-cytoplasmic ratio.

### FLIP quantification

For quantification of FLIP (Fluorescent Loss In Photobleaching) experiments, MetaMorph (Molecular Devices) was used to follow the integrated intensities of six different square ROIs of the same size as the bleached ROI: the bleached ROI, two reporting ROIs (one in the same compartment as the bleached point and one in the other compartment), two controls ROIs (one in the nucleus and one in the cytoplasm of one control cell) and one ROI outside any cells to measure the background intensity. The two nuclear ROIs and the cytoplasmic ROI of the bleached cells were positioned to be at approximately equal distances from one another. To normalize, first the background intensity was subtracted from the five other intensities measured in the cells. Then, the intensities measured in the cell of interest (three ROIs) were normalized by dividing by the average intensity of the two control ROIs. These data were plotted in the illustrative graphs.

To extract import and export rates, the nuclear and cytoplasmic compartments were determined manually in MATLAB using the roipoly command (Image Processing toolbox). The total population intensity of each compartment over time was then extracted based on these defined regions. Background subtraction and normalisation to the average of nuclear and cytoplasmic regions in a control cell was then carried out (as above). A simpler differential equation model than that employed in (Ege et al., 2018) was then fitted to this dynamic intensity data, including only temporal data and not spatial data. We fit to the population intensities using a system of ODEs given by

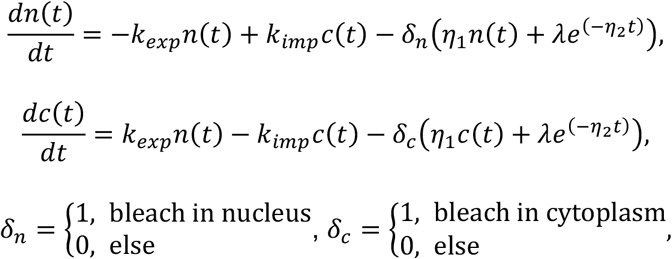

where *n*(*t*) and *c*(*t*) give, respectively, the total nuclear and cytoplasmic intensities at time, *t*. The parameter *k_exp_* gives the export rate from the nucleus to the cytoplasm and *k_imp_* the import rate from the cytoplasm to the nucleus. The final functions correspond to intensity decay due to bleaching depending on whether the bleach is in the nucleus or cytoplasm. A single decay parameter *η*_1_ allows the model to fit the bleach reasonably well, but underestimates the rapid initial bleach. Hence a decaying exponential function is also incorporated such that the bleach is effectively modelled by a double exponential, one with a rapid rate and the other a slower rate. This double exponential will clearly estimate the rate of bleaching better than a single exponential could (due to a greater degree of fitting freedom) and allows more robust estimates of population intensity transfer between compartments. The initial conditions are given by *n*(0) = *n*_0_ and *c*(0) = *c*_0_. At steady-state, prior to bleaching,

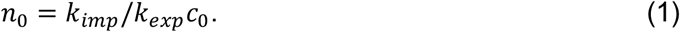

An analytic solution to the ODEs exists, however we estimated parameters by fitting a numerical solution of the model to the experimental intensity data for each cell. Parameter estimates for *k_exp_*, *k_imp_*, *n*_0_, *c*_0_, *η*_1_, *η*_2_ and *λ* were all required. We can reduce the number of parameters to fit by one by taking advantage of the steady-state relationship (1) to fix the initial nuclear intensity in terms of rates and initial cytoplasmic intensity. To optimise parameter fitting, we used the nonlinear model fitting function nlinfit in MATLAB’s Statistics and Machine Learning toolbox. Initial guesses for parameters in the nlinfit function were selected by choosing the initial cytoplasmic intensity in the experimental data for *c*_0_ and initial guess of order 10^−3^ based on population intensity analysis carried out in (Ege et al., 2018) for *k_exp_* and *k_imp_*. To generate initial guesses for the bleaching decay rates we fitted a double exponential function

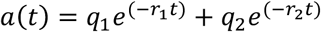

to the total intensity of the entire cell (nucleus and cytoplasm). Initial guesses to this double exponential fit were given as: 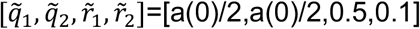. By differentiating this double exponential, *a*(*t*), into the form

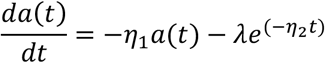

the output parameters fitted to the double exponential could be used as initial guesses for *η*_1_, *η*_2_ and *λ*.

### Optogenetic quantification

#### Opto_analyser – A Cell Compartment Segmentation App

To segment cellular compartments and extract dynamic intensities of the mitochondria, cytoplasm and nucleus we developed an app in MATLAB. The labelling of the nucleus and mitochondria alongside the GFP labelling of the protein of interest allow temporal segmentation of each compartment. This is done by the user selecting appropriate threshold levels in addition to carrying out a number of other processing steps. Appropriate compartment threshold levels differ from frame to frame and it would be too time consuming to threshold every compartment for every frame for every cell. In addition, in some frames the signal from a given compartment could be too weak to segment independently. Therefore, the movie is split into time windows of multiple frames. For example, for a movie of 100 frames, a window split of 4 would split the movie into four windows of 25 frames each whilst a single window split would provide only a single window covering all 100 frames. Intensity projections of each of these multiple framed windows are used in order to segment the compartments. A window split of 4 would then provide four segmented binary images that covered all 25 frames in each of their respective windows. This doesn’t provide total sensitivity to dynamic morphological changes but captures some of it whilst still being robust to high levels of noise and low levels of signal. A single window-split is obviously most robust to high noise and low signal but offers no sensitivity to morphological changes. Increasing the number of time windows leads to greater sensitivity to dynamic compartment movement within the movie but reduces robustness in segmentation due to the projections being taken from fewer frames in each window. In the analysis carried out here, a single percentile intensity projection of the entire movie (single window split) and percentile intensity projections of the movie split into 2, 4 and 8 time windows (multiple window splits) were considered.

The app was developed in MATLAB, using its inbuilt App Designer software and requires the Image Processing toolbox and lsmread from https://github.com/joe-of-all-trades/lsmread. The user interface is composed of three main panels. The left-hand side panel is for user input of file locations, requested number of window splits and save location. It also outputs error messages to guide the user. The central panel is composed of multiple tabs where the majority of image processing takes place. Each of these separate aspects of the image segmentation is carried out in a separate tab. Segmentation steps of the nuclear and mitochondrial compartments rely on the nuclear and mitochondrial channels only. Segmentation of the whole cell boundary relies on the generation of a hybrid channel where each frame is the maximum intensity projection of the nuclear, mitochondrial and protein channels in that frame. This makes the segmentation more robust to the heterogeneous and potentially weak intensity of the protein channel. Previews of the first frames in each channel allow the user to define each channel before zooming in on the region of interest around the cell. The app then iterates through each of the chosen time window splits in ascending order of number of windows (e.g. a single split is analysed first, followed by a two window split). For each selected window split, the app allows the user to define the intensity projections of the nuclear, mitochondrial and hybrid channels for all windows in that split simultaneously. The projections are generated via user defined percentiles of intensity of all frames in the relevant channel in that window. High percentiles allow objects to be visible for weak signals. Low percentiles can reduce noise and interference from neighbouring cells when the signals are strong. The optimum selected percentile is one that maximises the features of the cell under consideration but minimises interference from neighbouring cells and noise. Segmentation is carried out on these percentile intensity projections. Having selected the percentiles for all windows in that window split, the app then iterates through each of the window split’s windows. For each time window, the user manually crops around the cell to isolate the cell under consideration from neighbours, selecting from a choice of projections that best aid visual cropping. Using input sliders, they then select the desired threshold for segmentation of the hybrid channel, in order to determine the cell boundary in that time window. Images illustrating the current cell boundary are updated in real time as the threshold is adjusted. Having segmented the cell boundary, the user selects the most appropriate threshold levels for segmenting the nuclear and mitochondrial compartments. Again, images illustrating the compartment boundaries are updated in real time as the threshold levels are adjusted. Population intensities for each compartment over time are taken by using the segmented compartments as masks and applying them to the protein channel. The same compartment mask is used for all frames specific to each window. The right-hand side panel includes plots of each compartment population versus time, that update instantly according to chosen threshold levels, in addition to a record of all percentile and threshold choices for each time window in each window split. All previous population intensity profiles for previously completed window splits are included with easy to interpret colour mapping identifying each window split. For the current window split, the population intensity profiles include all windows up to and including the window currently being segmented. These plots and this data allows the user to determine the most appropriate thresholds not only from the thresholded images, but also from the emerging shape of the population intensity profiles and previously selected percentile and threshold levels. The presentation of all previous window-splits allows the user to investigate how the number of windows affects the resulting population intensity profiles. The plotting of all completed windows in the current window split allows the user to observe how smoothly the population intensity profile of the current window links with the previous one as the threshold levels of the current window are altered. Finally, once all window splits have been completed the segmentations are interpolated between time windows. A moving median filter is applied to intensity values of each frame of each compartment channel (nucleus, mitochondria and hybrid) with the size of the median filter being determined by the length of the windows for each split (i.e. if a movie of 100 frames is split into four windows of 25 frames each then the moving median filter has a range of 25 frames). Compartment thresholds are interpolated from the time window thresholds across all frames. Finally, the rough cell boundary crop (to limit interference from neighbouring objects) for each time window is interpolated across all frames by using distance transforms to model the change in shape from one binary crop to the next over the appropriate number of frames in the time window. This generates segmented compartments that change on a frame-by-frame basis rather than a window-by-window basis. Population intensities over time are then generated for these interpolated segmented compartments.

#### Background subtraction, bleaching intensity normalisation and non-conserved intensity correction methods

Intensity normalisation consists of three stages: i) background subtraction, ii) normalisation due to bleaching and iii) correcting for escape and return of the protein from the focal plane. The normalisation to account for bleaching should lead to the total intensity being constant over time and this is also what is assumed by the system of ODEs. However, upon release from the mitochondria, the protein of interest does not always remain in the viewing plane and instead escapes elsewhere in the cell. Similarly, during the recovery phase there can be a dramatic inflow of the protein of interest in the viewing plane.

Background subtraction is carried out for each frame. The mean of a 20 pixel by 20 pixel square in the top left hand corner of the movie is calculated over time and subtracted from the compartment intensities.

The intensities are then normalised to account for laser bleaching. The level of laser bleaching is determined from the behaviour of the total intensity of the cell over time, which appears to be nonlinear. The rate of bleaching appears to be composed of two approximately linear phases with gradients being approximately linear in both the release and recovery phases but with differing rates. As such, the total intensity is split into release and recovery phases and a linear fit made to each section. For all time prior to recovery, the bleach is assumed to fit the linear fit to total intensity in the release phase. From the first point of recovery, bleaching is assumed to fit the linear fit to bleaching of total intensity in the recovery phase.

Since the total intensity is not necessarily conserved due to inflow and outflow from the plane of view, the linear fits in each release and recovery phase are fit to temporal subsections. This avoids fitting a linear model to total intensity profiles that also include rapid outflow or inflow into the viewing plane in addition to low level bleaching. Let *C* be the median total intensity of the cell over the interval [0, *t_SS_end__*] where *t_SS_end__* is the last time-point prior to optorelease. Let *t*_*rl*_1__ be the beginning of the release fitting subsection and *t*_*rl*_2__ be the end. The linear fit to total intensity between *t*_*rl*_1__ and *t*_*rl*_2__ is given by 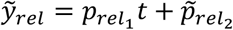 where we impose *p*_*rel*_1__ ≤ 0. Let *t*_*rc*_1__ be the beginning of the recovery fitting subsection and *t*_*rc*_2__ be the end. The linear total intensity fit between *t*_*rc*_1__ and *t*_*rc*_2__ is given by 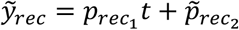 where we impose *p*_*rec*_1__ ≤ 0. To normalise to bleaching over these two phases requires a smooth bilinear fit to the gradual drop in total intensity. In the absence of inflow and outflow from the focal plane, these two linear fits, 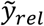 and 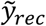 should smoothly connect at the end of the release and beginning of the recovery phase (*t* = *t*_*REC*_1__). The linear fit 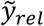 should also smoothly link with the median steady-state intensity, *C*, at timepoint *t_SS_end__* and linearly increase beyond this value for *t* < *t_SS_end__*. However, in cases where the total intensity is not conserved throughout the experiment, we must translate the linear functions 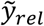 and 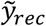 such that 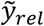 smoothly connects with *C* and 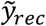. We adjust the y-intercept for the release phase according to

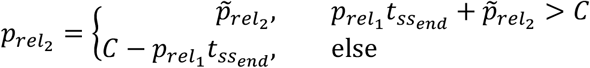

and define our translated linear fit as *y_rel_* = *p*_*rel*_1__ *t* + *p*_*rel*_2__. This links the release fit to the median steady-state intensity. We then adjust the y-intercept of 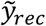 so it lines up with *y_rel_* by setting *p*_*rec*_2__ = (*p*_*rel*_1__ – *p*_*rec*_1__)*t*_*REC*_1__ + *p*_*rel*_2__. We then define our translated linear fit to the recovery phase as *y_rec_* = *p*_*rec*_1__ *t* + *p*_*rec*_2__. This gives us a smooth bilinear fit to drop in intensity due to bleaching that is independent of inflow and outflow and fit to differing bleach rates in the release and recovery phases.

Following background subtraction and normalisation of intensity to account for bleaching, we address the issue of non-conserved intensity due to rapid inflow and outflow from the image plane. In some cells, intensity is relatively well conserved. However, in a large number of cases we observe a rapid outflow from the image plane during release and rapid inflow during recovery. In a very small number of cases, we observe rapid inflow to the image plane during release and rapid outflow from the image plane during recovery. Due to the low number of cases of the latter, we only correct for the outflow-inflow case.

To detect cases of rapid outflow in release and rapid inflow in recovery we make use of the initial non-translated linear fits 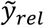 and 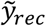 and the median steady-state intensity, *C*. Firstly, if 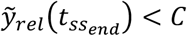 it suggests a discontinuous drop in total intensity upon release. This implies a rapid outflow from the focal plane upon release. Secondly, if 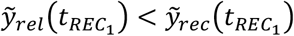 it suggests discontinuous increase in total intensity upon recovery. This implies a rapid inflow into the focal plane upon recovery. If both of these conditions are satisfied it suggests rapid outflow followed by rapid inflow that we can correct for in the model. If these conditions are satisfied, the narrow time periods of rapid outflow and inflow are identified and the total intensity over these time periods smoothed independently using Lowess. By considering the intensity profiles and the first derivative profiles, a smooth, monotonically decreasing outflow and smooth monotonically increasing inflow can be generated in the system. A vector of inflow and outflow with the smoothed intensity values over these two time windows and NaNs (not a number) everywhere else is generated.

#### Mathematical model

As in the FLIP analysis, we fit a system of ODEs to the extracted population intensity data for the mitochondrial, cytoplasmic and nuclear compartments. The ODEs are given by

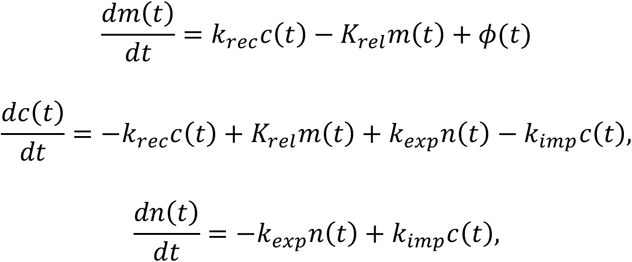

where *m*(*t*), *c*(*t*) and *n*(*t*) are the mitochondrial, cytoplasmic and nuclear intensities respectively at time, *t*. The initial conditions are given by *m*(0) = *m*_0_, *c*(0) = *c*_0_ and *n*(0) = *n*_0_. rate of export from the nucleus to the cytoplasm and rate of import from the cytoplasm to the nucleus are given by *k_exp_* and *k_imp_* respectively. The parameter *k_rec_* gives the rate at which the protein leaves the cytoplasm and binds to the mitochondria. The parameter *K_rel_* gives the rate at which protein leaves the mitochondria and enters the cytoplasmic compartment. The value *K_rel_* is dependent on whether the system is in the illumination phase or not i.e.

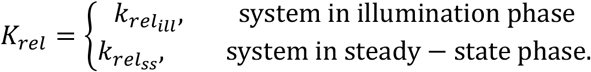

The final component of the ODEs is the function, *ϕ*(*t*), which accounts for cases where the total protein intensity in the system is not conserved in the experimental data, even after normalisation. We shall refer to this further in the model fitting methodology.

#### Model Fitting

The system of ODEs is fitted to the experimental data using MATLAB’s nlinfit function from the Statistics and Machine Learning toolbox. The fitting weights of each compartment are normalised such that each compartment contributes an equal amount to the residuals. Parameters that need fitting in the system are the rates *k_exp_*, *k_imp_*, *k_rec_*, *k_rel_ill__* and *k_rel_ss__* and the initial conditions *m*_0_, *c*_0_ and *n*_0_. The model is fit such that the parameter *k_rel_ill__* is fit only at the release phase and *k_rel_ss__* only fit at the initial steady-state phase and the recovery phase. The initial conditions need to be fitted since we cannot fix them at their experimental measurements due to noise in the system. We can reduce the number of parameters that need fitting by taking advantage of the steady-state relationships *c*_0_ = *k_rel_ss__/k_rec_m*_0_ and *n*_0_ = *k_imp_/k_exp_c*_0_ allowing us to fix the initial nuclear and cytoplasmic intensities in terms of other parameters. If the algorithm has identified that total intensity is conserved throughout the experiment then the inflow-outflow function *ϕ*(*t*) is set to zero everywhere. If the intensity is not conserved then *ϕ*(*t*) is constructed from the derivative of the inflow-outflow vector (described in the normalisation section) in the sections of smooth, monotonically decreasing release and smooth, monotonically increasing recovery. Everywhere else, where the inflow-outflow vector is composed of NaNs, the function *ϕ*(*t*) is set to zero. As initial guesses for our parameters we take 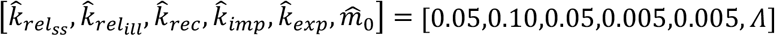 where *Λ* gives the median mitochondrial intensity over the first five frames prior to release.

For each cell, the intensity extraction was carried out by splitting each movie into 1, 2, 4 or 8 sections as described in the segmentation section. The system of ODEs was fit to each of these intensity extractions and the mean sum of squares of error for both the entire system and the fit to the nucleus only calculated. The fit with the lowest mean sum of squares of error was selected as being the optimum fit in each case.

## Acknowledgements

We thank members of the Crick Advanced Light Microscopy facility for technical support. We are indebted to and Hui Wang and Sirio Dupont for thoughtful discussion and advice.

## Author contributions

A.M.D. performed the experiments, R.P.J. developed the app and analytical methods, K.H. provided reagents prior to their publication, J.M.H. facilitated lattice light sheet imaging, T.F. and A.C. assisted with 3D data processing, M.M. and E.S. conceived the study, A.M.D., M.M., and E.S. wrote the manuscript.

## References

Aragona, M., Panciera, T., Manfrin, A., Giulitti, S., Michielin, F., Elvassore, N., Dupont, S. & Piccolo, S. 2013. A mechanical checkpoint controls multicellular growth through YAP/TAZ regulation by actin-processing factors. Cell, 154, 1047–1059.

Azzolin, L., Panciera, T., Soligo, S., Enzo, E., Bicciato, S., Dupont, S., Bresolin, S., Frasson, C., Basso, G., Guzzardo, V., Fassina, A., Cordenonsi, M. & Piccolo, S. 2014. YAP/TAZ incorporation in the beta-catenin destruction complex orchestrates the Wnt response. Cell, 158, 157–70.

Basu, S., Totty, N. F., Irwin, M. S., Sudol, M. & Downward, J. 2003. Akt phosphorylates the Yes-associated protein, YAP, to induce interaction with 14-3-3 and attenuation of p73-mediated apoptosis. Mol Cell, 11, 11–23.

Behar, M. & Hoffmann, A. 2010. Understanding the temporal codes of intra-cellular signals. Curr Opin Genet Dev, 20, 684–93.

Benham-Pyle, B. W., Pruitt, B. L. & Nelson, W. J. 2015. Cell adhesion. Mechanical strain induces E-cadherin-dependent Yap1 and beta-catenin activation to drive cell cycle entry. Science, 348, 1024–7.

Bertolio, R., Napoletano, F., Mano, M., Maurer-Stroh, S., Fantuz, M., Zannini, A., Bicciato, S., Sorrentino, G. & Del Sal, G. 2019. Sterol regulatory element binding protein 1 couples mechanical cues and lipid metabolism. Nat Commun, 10, 1326.

Beyer, H. M., Juillot, S., Herbst, K., Samodelov, S. L., Muller, K., Schamel, W. W., Romer, W., Schafer, E., Nagy, F., Strahle, U., Weber, W. & Zurbriggen, M. D. 2015. Red Light-Regulated Reversible Nuclear Localization of Proteins in Mammalian Cells and Zebrafish. ACS Synth Biol, 4, 951–8.

Cai, D., Feliciano, D., Dong, P., Flores, E., Gruebele, M., Porat-Shliom, N., Sukenik, S., Liu, Z. & Lippincott-Schwartz, J. 2019. Phase separation of YAP reorganizes genome topology for long-term YAP target gene expression. Nat Cell Biol, 21, 1578–1589.

Calvo, F., Ege, N., Grande-Garcia, A., Hooper, S., Jenkins, R. P., Chaudhry, S. I., Harrington, K., Williamson, P., Moeendarbary, E., Charras, G. & Sahai, E. 2013. Mechanotransduction and YAP-dependent matrix remodelling is required for the generation and maintenance of cancer-associated fibroblasts. Nat Cell Biol, 15, 637–46.

Chan, S. W., Lim, C. J., Loo, L. S., Chong, Y. F., Huang, C. & Hong, W. 2009. TEADs mediate nuclear retention of TAZ to promote oncogenic transformation. J Biol Chem, 284, 14347–58.

Chaudhuri, O., Gu, L., Klumpers, D., Darnell, M., Bencherif, S. A., Weaver, J. C., Huebsch, N., Lee, H. P., Lippens, E., Duda, G. N. & Mooney, D. J. 2016. Hydrogels with tunable stress relaxation regulate stem cell fate and activity. Nat Mater, 15, 326–34.

Chen, B. C., Legant, W. R., Wang, K., Shao, L., Milkie, D. E., Davidson, M. W., Janetopoulos, C., Wu, X. S., Hammer, J. A., 3RD, Liu, Z., English, B. P., Mimori-Kiyosue, Y., Romero, D. P., Ritter, A. T., Lippincott-Schwartz, J., Fritz-Laylin, L., Mullins, R. D., Mitchell, D. M., Bembenek, J. N., Reymann, A. C., Bohme, R., Grill, S. W., Wang, J. T., Seydoux, G., Tulu, U. S., Kiehart, D. P. & Betzig, E. 2014. Lattice light-sheet microscopy: imaging molecules to embryos at high spatiotemporal resolution. Science, 346, 1257998.

Codelia, V. A., Sun, G. & Irvine, K. D. 2014. Regulation of YAP by mechanical strain through Jnk and Hippo signaling. Curr Biol, 24, 2012–7.

Crefcoeur, R. P., Yin, R., Ulm, R. & Halazonetis, T. D. 2013. Ultraviolet-B-mediated induction of protein-protein interactions in mammalian cells. Nat Commun, 4, 1779.

De Mena, L., Rizk, P. & Rincon-Limas, D. E. 2018. Bringing Light to Transcription: The Optogenetics Repertoire. Front Genet, 9, 518.

Di Ventura, B. & Kuhlman, B. 2016. Go in! Go out! Inducible control of nuclear localization. Curr Opin Chem Biol, 34, 62–71.

Dupont, S., Morsut, L., Aragona, M., Enzo, E., Giulitti, S., Cordenonsi, M., Zanconato, F., Le Digabel, J., Forcato, M., Bicciato, S., Elvassore, N. & Piccolo, S. 2011. Role of YAP/TAZ in mechanotransduction. Nature, 474, 179–83.

Ege, N., Dowbaj, A. M., Jiang, M., Howell, M., Hooper, S., Foster, C., Jenkins, R. P. & Sahai, E. 2018. Quantitative Analysis Reveals that Actin and Src-Family Kinases Regulate Nuclear YAP1 and Its Export. Cell Syst, 6, 692–708 e13.

Elbediwy, A., Vincent-Mistiaen, Z. I., Spencer-Dene, B., Stone, R. K., Boeing, S., Wculek, S. K., Cordero, J., Tan, E. H., Ridgway, R., Brunton, V. G., Sahai, E., Gerhardt, H., Behrens, A., Malanchi, I., Sansom, O. J. & Thompson, B. J. 2016. Integrin signalling regulates YAP and TAZ to control skin homeostasis. Development, 143, 1674–87.

Elosegui-Artola, A., Andreu, I., Beedle, A. E. M., Lezamiz, A., Uroz, M., Kosmalska, A. J., Oria, R., Kechagia, J. Z., Rico-Lastres, P., Le Roux, A. L., Shanahan, C. M., Trepat, X., Navajas, D., Garcia-Manyes, S. & Roca-Cusachs, P. 2017. Force Triggers YAP Nuclear Entry by Regulating Transport across Nuclear Pores. Cell, 171, 1397–1410 e14.

Engelke, H., Chou, C., Uprety, R., Jess, P. & Deiters, A. 2014. Control of protein function through optochemical translocation. ACS Synth Biol, 3, 731–6.

Fang, L., Teng, H., Wang, Y., Liao, G., Weng, L., Li, Y., Wang, X., Jin, J., Jiao, C., Chen, L., Peng, X., Chen, J., Yang, Y., Fang, H., Han, D., Li, C., Jin, X., Zhang, S., Liu, Z., Liu, M., Wei, Q., Liao, L., Ge, X., Zhao, B., Zhou, D., Qin, H. L., Zhou, J. & Wang, P. 2018. SET1A-Mediated Mono-Methylation at K342 Regulates YAP Activation by Blocking Its Nuclear Export and Promotes Tumorigenesis. Cancer Cell, 34, 103–118 e9.

Fu, X., Liang, C., Li, F., Wang, L., Wu, X., Lu, A., Xiao, G. & Zhang, G. 2018. The Rules and Functions of Nucleocytoplasmic Shuttling Proteins. Int J Mol Sci, 19.

Gao, Z., Chen, S., Qin, S. & Tang, C. 2018. Network Motifs Capable of Decoding Transcription Factor Dynamics. Sci Rep, 8, 3594.

Guntas, G., Hallett, R. A., Zimmerman, S. P., Williams, T., Yumerefendi, H., Bear, J. E. & Kuhlman, B. 2015. Engineering an improved light-induced dimer (iLID) for controlling the localization and activity of signaling proteins. Proc Natl Acad Sci U S A, 112, 112–7.

Hao, N. & O’Shea, E. K. 2011. Signal-dependent dynamics of transcription factor translocation controls gene expression. Nat Struct Mol Biol, 19, 31–9.

Kanai, F., Marignani, P. A., Sarbassova, D., Yagi, R., Hall, R. A., Donowitz, M., Hisaminato, A., Fujiwara, T., Ito, Y., Cantley, L. C. & Yaffe, M. B. 2000. TAZ: a novel transcriptional co-activator regulated by interactions with 14-3-3 and PDZ domain proteins. EMBO J, 19, 6778–91.

Kennedy, M. J., Hughes, R. M., Peteya, L. A., Schwartz, J. W., Ehlers, M. D. & Tucker, C. L. 2010. Rapid blue-light-mediated induction of protein interactions in living cells. Nat Methods, 7, 973–5.

Kofler, M., Speight, P., Little, D., Di Ciano-Oliveira, C., Szaszi, K. & Kapus, A. 2018. Mediated nuclear import and export of TAZ and the underlying molecular requirements. Nat Commun, 9, 4966.

Lu, Y., Wu, T., Gutman, O., Lu, H., Zhou, Q., Henis, Y. I. & Luo, K. 2020. Phase separation of TAZ compartmentalizes the transcription machinery to promote gene expression. Nat Cell Biol, 22, 453–464.

Manning, S. A., Dent, L. G., Kondo, S., Zhao, Z. W., Plachta, N. & Harvey, K. F. 2018. Dynamic Fluctuations in Subcellular Localization of the Hippo Pathway Effector Yorkie In Vivo. Curr Biol, 28, 1651–1660 e4.

Meng, Z., Moroishi, T. & Guan, K. L. 2016. Mechanisms of Hippo pathway regulation. Genes Dev, 30, 1–17.

Molenaar, C. & Weeks, K. L. 2018. Nucleocytoplasmic shuttling: The ins and outs of quantitative imaging. Clin Exp Pharmacol Physiol, 45, 1087–1094.

Nakajima, H., Yamamoto, K., Agarwala, S., Terai, K., Fukui, H., Fukuhara, S., Ando, K., Miyazaki, T., Yokota, Y., Schmelzer, E., Belting, H. G., Affolter, M., Lecaudey, V. & Mochizuki, N. 2017. Flow-Dependent Endothelial YAP Regulation Contributes to Vessel Maintenance. Dev Cell, 40, 523–536 e6.

Niopek, D., Benzinger, D., Roensch, J., Draebing, T., Wehler, P., Eils, R. & Di Ventura, B. 2014. Engineering light-inducible nuclear localization signals for precise spatiotemporal control of protein dynamics in living cells. Nat Commun, 5, 4404.

Niopek, D., Wehler, P., Roensch, J., Eils, R. & Di Ventura, B. 2016. Optogenetic control of nuclear protein export. Nat Commun, 7, 10624.

Oka, T. & Sudol, M. 2009. Nuclear localization and pro-apoptotic signaling of YAP2 require intact PDZ-binding motif. Genes Cells, 14, 607–15.

Pocaterra, A., Romani, P. & Dupont, S. 2020. YAP/TAZ functions and their regulation at a glance. J Cell Sci, 133.

Purvis, J. E. & Lahav, G. 2013. Encoding and decoding cellular information through signaling dynamics. Cell, 152, 945–56.

Rosenbluh, J., Nijhawan, D., Cox, A. G., Li, X., Neal, J. T., Schafer, E. J., Zack, T. I., Wang, X., Tsherniak, A., Schinzel, A. C., Shao, D. D., Schumacher, S. E., Weir, B. A., Vazquez, F., Cowley, G. S., Root, D. E., Mesirov, J. P., Beroukhim, R., Kuo, C. J., Goessling, W. & Hahn, W. C. 2012. beta-Catenin-driven cancers require a YAP1 transcriptional complex for survival and tumorigenesis. Cell, 151, 1457–73.

Shreberk-Shaked, M. & Oren, M. 2019. New insights into YAP/TAZ nucleo-cytoplasmic shuttling: new cancer therapeutic opportunities? Mol Oncol, 13, 1335–1341.

Sorrentino, G., Ruggeri, N., Specchia, V., Cordenonsi, M., Mano, M., Dupont, S., Manfrin, A., Ingallina, E., Sommaggio, R., Piazza, S., Rosato, A., Piccolo, S. & Del Sal, G. 2014. Metabolic control of YAP and TAZ by the mevalonate pathway. Nat Cell Biol, 16, 357–66.

Strickland, D., Lin, Y., Wagner, E., Hope, C. M., Zayner, J., Antoniou, C., Sosnick, T. R., Weiss, E. L. & Glotzer, M. 2012. TULIPs: tunable, light-controlled interacting protein tags for cell biology. Nat Methods, 9, 379–84.

Strickland, D., Yao, X., Gawlak, G., Rosen, M. K., Gardner, K. H. & Sosnick, T. R. 2010. Rationally improving LOV domain-based photoswitches. Nat Methods, 7, 623–6.

Sudol, M. 2013. YAP1 oncogene and its eight isoforms. Oncogene, 32, 3922.

Tamm, C., Bower, N. & Anneren, C. 2011. Regulation of mouse embryonic stem cell self-renewal by a Yes-YAP-TEAD2 signaling pathway downstream of LIF. J Cell Sci, 124, 1136–44.

Valon, L., Marin-Llaurado, A., Wyatt, T., Charras, G. & Trepat, X. 2017. Optogenetic control of cellular forces and mechanotransduction. Nat Commun, 8, 14396.

Varelas, X., Miller, B. W., Sopko, R., Song, S., Gregorieff, A., Fellouse, F. A., Sakuma, R., Pawson, T., Hunziker, W., Mcneill, H., Wrana, J. L. & Attisano, L. 2010. The Hippo pathway regulates Wnt/beta-catenin signaling. Dev Cell, 18, 579–91.

Wada, K., Itoga, K., Okano, T., Yonemura, S. & Sasaki, H. 2011. Hippo pathway regulation by cell morphology and stress fibers. Development, 138, 3907–14.

Wang, H. & Hahn, K. M. 2016. LOVTRAP: A Versatile Method to Control Protein Function with Light. Curr Protoc Cell Biol, 73, 21 10 1–21 10 14.

Wang, H., Vilela, M., Winkler, A., Tarnawski, M., Schlichting, I., Yumerefendi, H., Kuhlman, B., Liu, R., Danuser, G. & Hahn, K. M. 2016a. LOVTRAP: an optogenetic system for photoinduced protein dissociation. Nat Methods, 13, 755–8.

Wang, S., Lu, Y., Yin, M. X., Wang, C., Wu, W., Li, J., Wu, W., Ge, L., Hu, L., Zhao, Y. & Zhang, L. 2016b. Importin alpha1 Mediates Yorkie Nuclear Import via an N-terminal Non-canonical Nuclear Localization Signal. J Biol Chem, 291, 7926–37.

Xu, L. & Massague, J. 2004. Nucleocytoplasmic shuttling of signal transducers. Nat Rev Mol Cell Biol, 5, 209–19.

Yosef, N. & Regev, A. 2011. Impulse control: temporal dynamics in gene transcription. Cell, 144, 886–96.

Yumerefendi, H., Dickinson, D. J., Wang, H., Zimmerman, S. P., Bear, J. E., Goldstein, B., Hahn, K. & Kuhlman, B. 2015. Control of Protein Activity and Cell Fate Specification via Light-Mediated Nuclear Translocation. PLoS One, 10, e0128443.

Yumerefendi, H., Lerner, A. M., Zimmerman, S. P., Hahn, K., Bear, J. E., Strahl, B. D. & Kuhlman, B. 2016. Light-induced nuclear export reveals rapid dynamics of epigenetic modifications. Nat Chem Biol, 12, 399–401.

Zanconato, F., Cordenonsi, M. & Piccolo, S. 2016. YAP/TAZ at the Roots of Cancer. Cancer Cell, 29, 783–803.

Zanconato, F., Cordenonsi, M. & Piccolo, S. 2019. YAP and TAZ: a signalling hub of the tumour microenvironment. Nat Rev Cancer, 19, 454–464.

Zhao, B., Li, L., Tumaneng, K., Wang, C. Y. & Guan, K. L. 2010. A coordinated phosphorylation by Lats and CK1 regulates YAP stability through SCF(beta-TRCP). Genes Dev, 24, 72–85.

